# Systematic CRISPRi perturbation of 1,408 autism risk genes maps multilevel transcriptional convergence in human cortical neurons

**DOI:** 10.64898/2026.09.02.748925

**Authors:** Ashlesha Gogate, Wei-Chen Chen, Mpathi Nzima, Chikara Takeuchi, Huan Zhao, Lei Wang, Sushama Sivakumar, Minnie Deng, Kiran Kaur, W. Lee Kraus, Nikhil V. Munshi, Gary C. Hon, Maria H. Chahrour

## Abstract

Genetic heterogeneity in autism spectrum disorder (ASD) complicates the identification of shared molecular pathways amenable to therapeutic intervention. Here, we perform a CRISPRi Perturb-seq screen targeting 1,408 ASD risk genes in human embryonic stem cell-derived immature cortical neurons and profile the resulting transcriptomic effects by single-cell RNA sequencing. We identify 215 ASD risk genes whose repression induces significant global transcriptomic dysregulation. Leveraging this functional atlas, we characterize candidate genes based on transcriptional similarity to established ASD hub genes. We identify gene programs recurrently dysregulated across perturbations, anchored by processes governing microtubule dynamics, neuron differentiation, cell migration, and transmembrane transport. We further identify CHAMP1 as a previously unrecognized regulator of Wnt signaling and define specific ASD risk genes that modulate the rate of cortical neuron differentiation. Analysis of differentially expressed genes reveals both convergent and perturbation-specific downstream transcriptional responses. Together, these findings provide a multilevel map of transcriptomic convergence in ASD and establish a framework for identifying both pathway-level and genotype-specific therapeutic strategies.

## Introduction

Autism spectrum disorder (ASD) is a collection of individually rare neurodevelopmental disorders with heterogeneous clinical presentations that converge on core features including impaired social communication, restricted interests, and repetitive behaviors^1,2^. ASD affects ∼1-1.5% of the global population^3^. Its heritability is estimated at ∼80-93%, yet no single genetic locus accounts for more than a small fraction of population-level risk^4^. The genetic architecture of ASD is highly complex, comprising both rare variants with large effect sizes (accounting for ∼15-20% of overall genetic risk) and common variants that individually contribute minimally to risk (collectively accounting for least 50% of ASD risk)^5^. Large-scale whole-exome and whole-genome sequencing studies have identified hundreds of ASD risk genes through rare *de novo* and inherited variant mapping^6–10^. Integrating these datasets, the Simons Foundation Autism Research Initiative (SFARI) Gene database now catalogues 1,267 ASD risk genes (2026 Q1 release), a number that continues to grow as additional sequencing cohorts are analyzed^11^.

Despite this remarkable genetic heterogeneity, convergent patterns have emerged at the transcriptomic level. Post-mortem ASD brain studies have consistently identified dysregulated gene co-expression networks and implicated specific developmental stages and cell types in ASD pathogenesis^12–17^. ASD risk genes are co-expressed in the developing brain and form protein-protein interaction networks, pointing to shared molecular architecture^13,18–20^. Consequently, the field has shifted toward identifying the downstream biological pathways through which genetically diverse risk factors converge, with the goal of identifying mechanisms amenable to therapeutic intervention. Studies using human stem cell-derived cultures, cerebral organoids, assembloids, and animal models have begun to map these convergent effects in parallel^19,21–34^, collectively demonstrating that genetically diverse and constrained risk factors functionally converge on fundamental processes including early cortical neurogenesis, neuronal migration, progenitor proliferation, chromatin regulation, and synaptic signaling.

However, existing functional screens remain limited in scale, experimental design, and resolution, restricting biological inference. Most studies examine relatively small, manually curated gene panels that represent only a fraction of the known ASD risk genes, introducing selection bias toward well-studied loci. The majority rely on complete gene knockouts, which do not recapitulate the haploinsufficiency underlying most ASD risk variants. Many employ low-content phenotypic readouts, such as cell survival or gross morphology, rather than comprehensive single-cell transcriptomic profiling. Systems such as cerebral organoids and assembloids, while biologically rich, introduce substantial phenotypic variability that complicates direct comparisons across perturbations. And while studies of adult post-mortem cortex provide valuable human tissue context, they cannot capture the early developmental stages during which ASD risk genes are most functionally relevant. No single study has simultaneously addressed all these limitations at the scale of the full catalogue of ASD risk genes.

Here, we perform a pooled CRISPR interference (CRISPRi) Perturb-seq screen targeting 1,408 ASD risk genes in human embryonic stem cell (hESC)-derived immature cortical neurons, with single-cell RNA sequencing (scRNA-seq) as the phenotypic readout. At the whole-transcriptome level, perturbation of 215 ASD risk genes induced significant transcriptional dysregulation, grouping into 31 functional clusters based on transcriptional similarity. We identified gene programs recurrently dysregulated across perturbations and organized perturbations into six major biological themes using mixture modeling. We validated CHAMP1 as a previously unrecognized regulator of Wnt signaling, identified ASD risk genes that modulate the rate of cortical neuron differentiation, and mapped differentially expressed coding genes, long non-coding RNAs (lncRNAs), and transposable elements across the full perturbation landscape. Together, these findings demonstrate multilevel transcriptomic convergence across genetically diverse ASD risk genes and provide a high-resolution functional atlas for investigating ASD biology and identifying therapeutic targets.

## Methods

### Human embryonic stem cell culture

H9 human embryonic stem cells (hESCs) were cultured as previously described^35^. Briefly, H9 cells were maintained under feeder-free conditions in mTeSR Plus medium (STEMCELL Technologies) and incubated at 37°C and 5% CO_2_. The H9 dCas9-Krab (H9 dCK) hESC line was used for Perturb-seq^35^.

### Neuronal differentiation

Neuronal differentiation was performed using a modified version of the protocol described by Qi et al., 2017^36^. Briefly, hESCs were seeded on Matrigel-coated plates or 12 mm coverslips at a density of 200,000 cells/cm^2^ or ∼80-90% confluence. The culture medium (mTeSR) was replaced with KSR medium, consisting of 17:3 knockout DMEM to 3 knockout serum replacement, 1 mM glutamine, 100 µM MEM non-essential amino acids, and 0.1 mM β-mercaptoethanol. On differentiation days 0 and 1 (D0, D1), the medium was supplemented with 250 nM LDN193189, 10 µM SB431542, and 5 µM XAV939. On D2 and D3, the medium was further supplemented with 8 µM PD0325901, 10 µM SU5402, and 10 µM DAPT. On D4 and D5, the medium was gradually transitioned by replacing one-third of the KSR medium with N2 medium supplemented with B27, while maintaining the composition of the D2-D3 medium. On D6 and D7, the KSR ratio was adjusted to 1:2, and LDN193189, SB431542, and XAV939 were omitted. Medium was changed every 24 hours from D0 through D7. On D8, immature cortical neurons were dissociated with Accutase for 30 minutes at 37°C and replated in neuronal medium consisting of Neurobasal medium supplemented with B27, 20 ng/ml BDNF, 0.5 mM dibutyryl cAMP, and 0.2 mM ascorbic acid, with PD0325901, SU5402, and DAPT. Cells were seeded onto 12 mm coverslips coated with poly-L-ornithine (15 µg/ml), mouse laminin I (1 µg/ml), and fibronectin (2 µg/ml) in PBS for 24 hours at 37°C at a density of 300,000 cells/cm^2^. The medium was completely replaced with fresh neuronal medium on D9 and D11, and every 3-4 days thereafter. On D20, neurons were sparsely transfected with a GFP-expressing plasmid (pCAG-EGFP) using Lipofectamine 2000 (Invitrogen). Conditioned medium was temporarily replaced with serum- and antibiotic-free medium containing DNA-Lipofectamine complexes (1 µg plasmid DNA per well) for 2 hours. The conditioned medium was then returned to the cultures for a 24-hour recovery period. Cells were collected at different time points for downstream analyses. For Perturb-seq and RNA extraction, D6 cells were dissociated with Accutase for 30 minutes at 37°C. For quality control, D6 cells grown on coverslips were fixed with 4% paraformaldehyde for 15 minutes at room temperature with shaking and stored in cold PBS for immunocytochemistry. For immunofluorescence analyses, D21 neurons were fixed and stored using the same procedure.

### Design of the sgRNA library

sgRNA sequences for most target genes were obtained from Replogle et al., 2022^37^. Six gRNAs were included for each target transcript. The final library also contained 600 non-targeting sgRNAs as negative controls and 600 targeting sgRNAs as positive controls. sgRNAs for the remaining targets were designed using FlashFry^38^.

### CRISPRi Perturb-seq

Library construction, lentiviral packaging, lentiviral transduction, and single-cell RNA sequencing (scRNA-seq) were performed as previously described^35^. Briefly, pooled DNA oligonucleotides (Twist Bioscience) were cloned into LentiGuide(10X)-BFP-Puro (Addgene #229014) vector and electroporated into Endura Duo competent cells (Lucigen, 60242-2). For lentiviral production (per 10 cm plate; 10 total plates), 8 µg of the sgRNA library plasmid was co-transfected into HEK293T cells with 2 µg of pMD2.G and 6 µg of psPAX2 packaging plasmids (Addgene #12259 and #12260, respectively), along with 64 µl Transporter 5 (Kyfora Bio, 26008) and 1 ml Opti-MEM. Culture medium was replaced 24 hours post-transfection, and viral supernatants were collected just under 48 hours later (just under 72 hours post-transfection). Harvested lentivirus was concentrated 30-fold by ultracentrifugation prior to transduction. H9 dCK hESCs were harvested at 80% confluency and transduced as a single-cell suspension. Following transduction, fluorescence-activated cell sorting (FACS) was used to select cells in the 40^th^-90^th^ percentile of BFP-positive cells to enrich for target multiplicity of infection (2-4 sgRNAs per cell), followed by expansion for 2 weeks before neuronal differentiation.

For scRNA-seq, cells were loaded onto a Chromium Next GEM Chip N (10x Genomics) using hash-tag oligonucleotide (HTO) superloading^39^. Raw sequencing data were processed according to the manufacturer’s instructions (10x Genomics User Guide CG000513).

### scRNA-seq data preprocessing and filtering

scRNA-seq data were preprocessed using an in-house pipeline, Perturb-Seq-Processing-Pipeline, available on GitHub (https://github.com/Hon-lab/Perturb-Seq-Processing-Pipeline/). Briefly, transcriptomic reads were aligned to the human reference genome (hg38) using Cell Ranger (v7.0.0, 10x Genomics). Cells with low UMI counts, greater than 20% mitochondrial transcripts, or poor-quality transcriptomes were excluded. Genes with fewer than one count across the dataset were removed. sgRNAs and HTO were assigned to cells using FBA^40^. Only singlet cells (i.e., cells assigned a single HTO) containing at least 1 sgRNA were retained for downstream analyses. Fifty principal components were used for UMAP embedding. Clonal expansion can confound Perturb-seq analyses by introducing transcriptional signatures associated with shared lineage rather than the targeted genetic perturbation^41^. To minimize these effects, clonally expanded cells identified by highly recurrent sgRNA combinations were collapsed by randomly retaining a single representative cell from each clone, while all remaining cells from the same clone were excluded from downstream analyses.

### Energy distance analysis

Energy distance analysis was performed as previously described^37,42,43^. Briefly, analyses were performed using the top 50 principal components (PCs). To determine whether two distinct cell populations exhibit significantly different transcriptomic profiles, we calculated energy distance defined as (2A - B - C), where (A) represents the average multivariate distance between two cell populations (Group X and Group Y), and (B) and (C) represent the within-group dispersion for Groups X and Y, respectively. Positive energy distance values indicate that between-group variation exceeds within-group variation, reflecting differences in the underlying transcriptomic distributions between populations.

To identify potential guide-specific outliers, distance components (DISCO), a non-parametric extension of ANOVA, was first applied to each target. Guide labels were randomly permuted among cells within each target group across independent iterations to generate a locus-specific null distribution. Empirical *P* values were calculated as the proportion of permuted statistics that were equal to or greater than the observed statistic. Targets with DISCO *P* > 0.05 were considered to lack evidence for guide-specific effects, and all guides were retained. Targets with DISCO *P* ≤ 0.05 were further evaluated for potential outlier guides. For targets failing the DISCO assessment, unsupervised k-means clustering (k = 2) was performed to identify a small, high-variance cell cluster distant from the dominant cluster centroid. Guides enriched within this cluster were flagged as potential outliers and removed. Following outlier guide removal, energy distance was calculated between cells receiving targeting sgRNAs and non-targeting control (NTC) sgRNAs. For every target, 2,000 cells receiving NTC sgRNAs were randomly sampled and 1,000 permutations were performed. This was repeated 20 times, amounting to 20,000 permutations per target. Targets were considered significant hits if they met a minimum energy distance cutoff of 0 and a significance threshold of *P* ≤ 0.001, calculated across 20 permutation batches.

Differences in the distribution of loss-of-function observed/expected upper bound fraction (LOEUF) scores between significant and non-significant hits were assessed using the Anderson-Darling k-sample test. A target-by-target distance matrix was generated from pairwise energy statistics across high-dimensional transcriptional profiles of significant perturbations. Perturbations were subsequently clustered using affinity propagation to identify unbiased groups of transcriptionally similar perturbations.

### Cell proportion analysis

To determine whether individual ASD risk gene perturbations significantly altered cell type proportions compared with NTC, we adapted the statistical framework developed for pooled single-cell CRISPR screens by Li et al., 2023^29^. Briefly, we performed a Cochran-Mantel-Haenszel (CMH) test stratified by sequencing library to evaluate the overall odds ratio for each perturbation-cluster pair. Raw *P* values were adjusted for multiple testing using the Benjamini-Hochberg method. To distinguish biological effects from technical variation, we constructed a cluster-specific empirical null distribution. We performed 100 permutations in which cell barcode labels were randomly shuffled relative to the sgRNA count matrix, and pseudo-target perturbations were generated by randomly sampling sets of 6 sgRNAs. For each permutation, stratified CMH tests were recalculated to generate cluster-specific empirical null distributions. Observed log odds ratios were standardized relative to the corresponding cluster-specific null distribution to compute an empirical t statistic. Empirical *P* values were then calculated from these distributions and adjusted for multiple testing using the Benjamini-Hochberg method. Perturbations with both a multiple-testing-adjusted CMH *P* < 0.05 and a multiple-testing-adjusted empirical *P* < 0.05 were considered to significantly alter cell proportions.

### Pseudotime analysis

Monocle3 (v1.3.4)^44,45^ was used for pseudotime analysis. A cell dataset object was generated using normalized RNA expression counts, dimensionality reduction coordinates, and clustering information. All cells were assigned to a single partition. The learn_graph function was used to construct a principal graph, and cells were interactively ordered along the trajectory based on their progression from less differentiated to more differentiated states. Neural progenitor cells were selected as the trajectory root to define the starting point of pseudotime ordering. Trajectories were inferred in reduced-dimensional space using reversed graph embedding. Pseudotime values were scaled from 0 to 1 for visualization.

### Generation of individual knockdown cell lines

The sgRNAs that were most effective at repressing the target genes were selected from the Perturb-seq screen. These sgRNAs were individually cloned into the PB9.1 plasmid^35^, sequence-verified, and nucleofected into H9 dCK hESCs. Typically, two sgRNAs targeting the same gene were nucleofected simultaneously to ensure efficient repression of the target gene. Four days after nucleofection, cells were treated with 1 µg/ml puromycin for 1 week to select for sgRNA-containing hESCs. After selection, cells were allowed to recover for 7-10 days, after which target gene repression was assessed by qRT-PCR.

### Immunohistochemistry

Immunostaining was performed for differentiation quality control and to assess the effects of perturbations on neuronal development. Cells were grown on coverslips, fixed, and permeabilized with 0.5% Triton X-100 in PBS (PBS-T) for 5 minutes at room temperature. Coverslips were blocked with blocking buffer (5% normal goat serum in 0.1% PBS-T) for 1 hour at room temperature and subsequently incubated overnight at 4°C with primary antibodies diluted in blocking buffer. The primary antibodies used were OCT3/4 (Sata Cruz, sc-5279, 1:100), DCX (Invitrogen, 48-1200, 1:200), GFP (Abcam, 1018753-39, 1:500), and TUJ1 (Covance, MRB-435P, 1:500). Following primary antibody incubation, coverslips were washed and incubated with secondary antibodies (Alexa Fluor 488, Alexa Fluor 555, and Alexa Fluor 647; Invitrogen) diluted 1:1000 in blocking buffer for 1 hour at room temperature. Coverslips were mounted onto slides using antifade mounting medium containing DAPI (Vector Laboratories). Immunofluorescence images were acquired using a Zeiss LSM880 confocal microscope with a 20X objective. Images were processed and analyzed using the Simple Neurite Tracer plugin (SNT, v4.2.1) in Fiji (ImageJ v2.14.0/1.54f). Data were assessed for normality using the Shapiro-Wilk test to determine the appropriateness of parametric testing. An F-test was used to assess equality of variances and to determine whether an unpaired t-test or Welch’s t-test was appropriate. Statistical analyses were performed using GraphPad Prism.

### Gene program analysis

To identify gene programs (GPs), consensus non-negative matrix factorization (cNMF)^46^ was applied to the raw count matrix from the filtered dataset. Factorization was performed for 20 iterations across the top 2,000 highly variable genes using K values of 50, 100, 200, 250, and 500. A local density threshold of 0.1 was used during consensus clustering to consolidate program matrices across iterations^42^, generating a cell-by-gene program usage matrix and a gene spectra score matrix for each K value.

For each GP, driver genes were identified using two approaches: (1) the top 300 genes ranked by gene spectra score within each GP and (2) a knee detection algorithm^47^ applied to the ranked gene spectra score distribution to identify the inflection point. Gene ontology (GO) enrichment analysis was performed per GP independently for both driver gene sets. To determine whether individual perturbations significantly dysregulated specific GPs, mean GP usage scores were calculated for each perturbation and compared with NTC cells. Log_2_ fold changes were calculated, and statistical significance was assessed using the Mann-Whitney U test. *P* values were adjusted using the Benjamini-Hochberg false discovery rate (FDR) correction, with FDR < 0.001 used as the significance threshold. The optimal number of GPs (K = 200) was selected based on the Forbenius error curve, the number of unique GO terms recovered, the number of unique significant perturbations identified, and cosine similarity between GPs.

The top 300 driver genes for each GP were overlapped with autism spectrum disorder (ASD) risk genes from the SFARI Gene database (2025 Q3 release). Statistical significance of overlap was assessed using a hypergeometric test followed by FDR correction. GPs with adjusted *P* < 0.05 were considered significantly enriched for ASD risk genes. To evaluate the developmental relevance and cell type-specific expression of identified *in vitro* GPs, we analyzed a published snRNA-seq dataset spanning prenatal and postnatal human cortex development^48^ and a spatial transcriptomics dataset derived from the adult human dorsolateral prefrontal cortex^49^. Donor specimens were grouped into four developmental windows: fetal (second and third trimester), early postnatal (0-4 years), late postnatal (4-20 years), and adult. GP usage scores were calculated for each cell in the snRNA-seq and the spatial transcriptomics datasets using the scanpy.tl.score_genes function^50^ and the UCell library^51^, respectively, based on the top 300 genes ranked by gene spectrum score for each GP.

To classify perturbations into higher-order functional groups based on GP dysregulation profiles, a latent class mixture modeling framework was implemented using the StepMix library^52^. Only targeted gene perturbations represented by ≥ 100 cells were included. For each perturbation, the NTC baseline vector (mean normalized GP usage across NTC cells) was subtracted from the mean normalized GP usage vector, and each GP was subsequently z-scored to mean of 0 and variance of 1. The number of cells per perturbation was included as a covariate in the model. Finite Gaussian mixture models containing 2-15 latent classes were evaluated, with 200 independent model fits performed for each class number using a single initialization strategy across multiple random seeds. Model selection was based on minimization of Bayesian information criterion (BIC) and Akaike information criterion (AIC), identifying 5 latent classes as the optimal solution. Following class number selection, the final model was refit 200 times using multiple independent initializations and a single random seed to ensure model convergence. Latent class assignments were extracted from the final posterior probability matrix. Mean z-scored GP usage values were calculated for each class for visualization.

### qRT-PCR

Total RNA was extracted from hESCs and day 6 neuronal cultures. Superscript III First-Strand Synthesis System (Invitrogen) was used to reverse transcribed 2 µg of RNA into cDNA. Quantitative real-time PCR was performed using PowerUp SYBR Green Master Mix (2X; Applied Biosystems) and the gene-specific primers listed below.

*ASCL1* forward: 5’-CCCAAGCAAGTCAAGGCACA-3’, reverse: 5’-AAGCCGCTGAAGTTGAGCC-3’;

*CHAMP1* forward: 5’-CGTTCCCTGCTGTCTCCCCAGA-3’, reverse: 5’-GTGGCCCTGGTTTCCAAGAGCC-3’;

*DCX* forward: 5’-TATGCGCCGAAGCAAGTCTCCA-3’, reverse: 5’-CATCCAAGGACAGAGGCAGGTA-3’;

*GABRG2* forward: 5’-GTCTGGACGGCAAGGACTGTGC-3’, reverse: 5’-ACAGGCAGAAGGCAGTGGGGAA-3’;

*GAPDH* forward: 5’-CCAGCGAGATCCCTCCAAAAT-3’, reverse: 5’-GGCTGTTGTCATACTTCTCATGG-3’;

*GLI3* forward: 5’-TCATGGAGGCCCAGTCCCACAG-3’, reverse: 5’-AGGCAACGGCTTTCTCGCTCAC-3’;

*GREM1* forward: 5’-AGGCTGCAACAGTCGCACCATC-3’, reverse: 5’-TTCCGGATGTGCCTGGGGATGT-3’;

*ISL1* forward: 5’-CGTGCCCGCTCCAAGGTGTATC-3’, reverse: 5’-AAGCGCAAATTCGTCCCCAGGG-3’;

*MXRA8* forward: 5’-ACACCCCTCCCTTGGACTCTGC-3’, reverse: 5’-GTGGGGTGCTGAGAGTGGCAAC-3’;

*OCT4* forward: 5’-CCTGAAGCAGAAGAGGATCACC -3’, reverse: 5’-AAAGCGGCAGATGGTCGTTTGG-3’;

*PAX3* forward: 5’-CACCGTGCCGTCAGTGAGTTCC-3’, reverse: 5’-CCGTCGATGCTGTGTTTGGCCT-3’;

*PAX6* forward: 5’-AACGATAACATACCAAGCGTGT-3’, reverse: 5’-GGTCTGCCCGTTCAACATC-3’;

*PITX2* forward: 5’-TCCCATGTCTTCGTTTGCCCGC-3’, reverse: 5’-CCCGCCGAGTTCTCAAGCCAAG-3’;

*RPGRIP1L* forward: 5’-GCACGCTAGGCCATGTCTGGTC-3’, reverse: 5’-CTGACACGTGACACTGCCTGGC-3’;

*TBR1* forward: 5’-TCACTGGAGGTTTCAAGGAGGC-3’, reverse: 5’-AAGCCGCTGAAGTTGAGCC-3’;

*TCF7L2* forward: 5’-GCAGCACCCTCACCATGTCCAC-3’, reverse: 5’-ATATCTGGAGGGTGCGGAGGCC-3’;

*WNT1* forward: 5’-TGGGTTTCTGCTACGCTGCTGC-3’, reverse: 5’-AAGCAGGTTCGTGGAGGAGGCT-3’.

### Differentially expressed gene (DEG) analysis

pySpade (v0.1.7)^53^ was used to identify differentially expressed genes (DEGs) for each perturbation. The process command was used to reformat transcriptomic and sgRNA assignment outputs from the in-house preprocessing pipeline for downstream analysis. The DEobs function performed differential expression analysis by comparing each targeted perturbation with NTC cells. The DErand function performed differential expression analysis using randomly selected cell populations of varying sizes (n = 100, 200, 300, 400, 500, 600, 700, 800, 900, 1,000, 1,250, 1,500, 1,750, 2,000, and 2,500) compared with NTC cells, with randomization set to “equal”. The global function calculated DEG significance scores by integrating observed and randomized *P* values using a gamma distribution approximation. To calibrate the significance threshold, pySpade was also applied to a shuffled sgRNA assignment matrix, resulting in a significance score cutoff of - 8.5 corresponding to an FDR < 0.1. Reported DEG hits met the criteria of FDR < 0.1, absolute fold change > 20%, and expression in at least 5% of cells.

### Long non-coding RNA (lncRNA) analysis

Genomic coordinates for human lncRNAs were downloaded from BioMart^54^, converted to genomic ranges, and annotated with the nearest protein-coding gene (PCG) based on genomic proximity. DEGs identified for each perturbation were classified as either lncRNAs or PCGs. A putative cis-regulatory relationship was defined when a lncRNA and its nearest PCG were both independently identified as DEGs within the same perturbation. Co-dysregulated lncRNA-PCG pairs were overlapped with ASD risk genes from the SFARI Gene database (2025 Q3 release) to annotate PCGs associated with ASD risk. lncRNAs from identified co-dysregulated pairs were further annotated using three regulatory element datasets: (1) chromatin state segmentation data from nine non-neuronal human cell lines (ENCODE)^55^, (2) H3K4me3 profiles from human prefrontal cortex samples from 11 individuals (uMass)^56^, and (3) predicted developmental brain enhancers derived from fetal brain samples (CBA)^57^. lncRNAs overlapping H3K4me3 peaks in the uMass dataset were classified as “predicted human brain promoter” lncRNAs. Among these, lncRNAs that did not overlap regions annotated as “1_Active_Promoter” in any of the nine ENCODE non-neuronal cell lines were further classified as “predicted human brain-specific promoter” lncRNAs. lncRNAs overlapping predicted regulatory elements in the CBA dataset were classified as “predicted human brain enhancer” lncRNAs. Among these, lncRNAs that did not overlap regions annotated as “4_Strong_Enhancer” or “5_Strong_Enhancer” in any of the nine ENCODE non-neuronal cell lines were further classified as “predicted human brain-specific enhancer” lncRNAs.

### Transposable element (TE) analysis

TE expression was quantified from alignment files using the scTE pipeline^58^ with genomic-exclusive reference indices generated from the hg38 genome assembly. Approximately 75,000 cells per library were specified as the expected cell number for quantification. Individual TE expression matrices were merged into a single matrix, and differentially expressed TEs were identified using pySpade^53^ as described above. TE family and phylogenetic annotations were obtained from Dfam 3.3^59^.

## Results

### Perturbation of 1,408 ASD risk genes in hESC-derived cortical neurons prioritizes 215 genes that significantly alter the transcriptome

To investigate the role of autism spectrum disorder (ASD) risk genes in early cortical development, we curated a gene list from the Simons Foundation Autism Research Initiative (SFARI) Gene database (2025 Q3 release)^11^, large-scale ASD genetic studies^6–9^, genes located within ASD-associated copy number variant (CNV) loci (16p11.2 syndrome, DiGeorge syndrome, and Williams syndrome), and additional candidate genes^60–62^ (Figure 1a). Many genes initially nominated by large-scale sequencing studies have since been catalogued in SFARI, reflecting ongoing convergence across independent gene discovery efforts. We performed CRISPRi Perturb-seq targeting 1,408 ASD risk genes in human embryonic stem cell (hESC)-derived immature cortical neurons (Figure 1b, Supplementary Figure S1, Supplementary Table S1). We confirmed efficient on-target repression (average repression 58%) and reproducibility across replicate libraries (Supplementary Figure S2, Supplementary Table S2).

**Figure 1.**
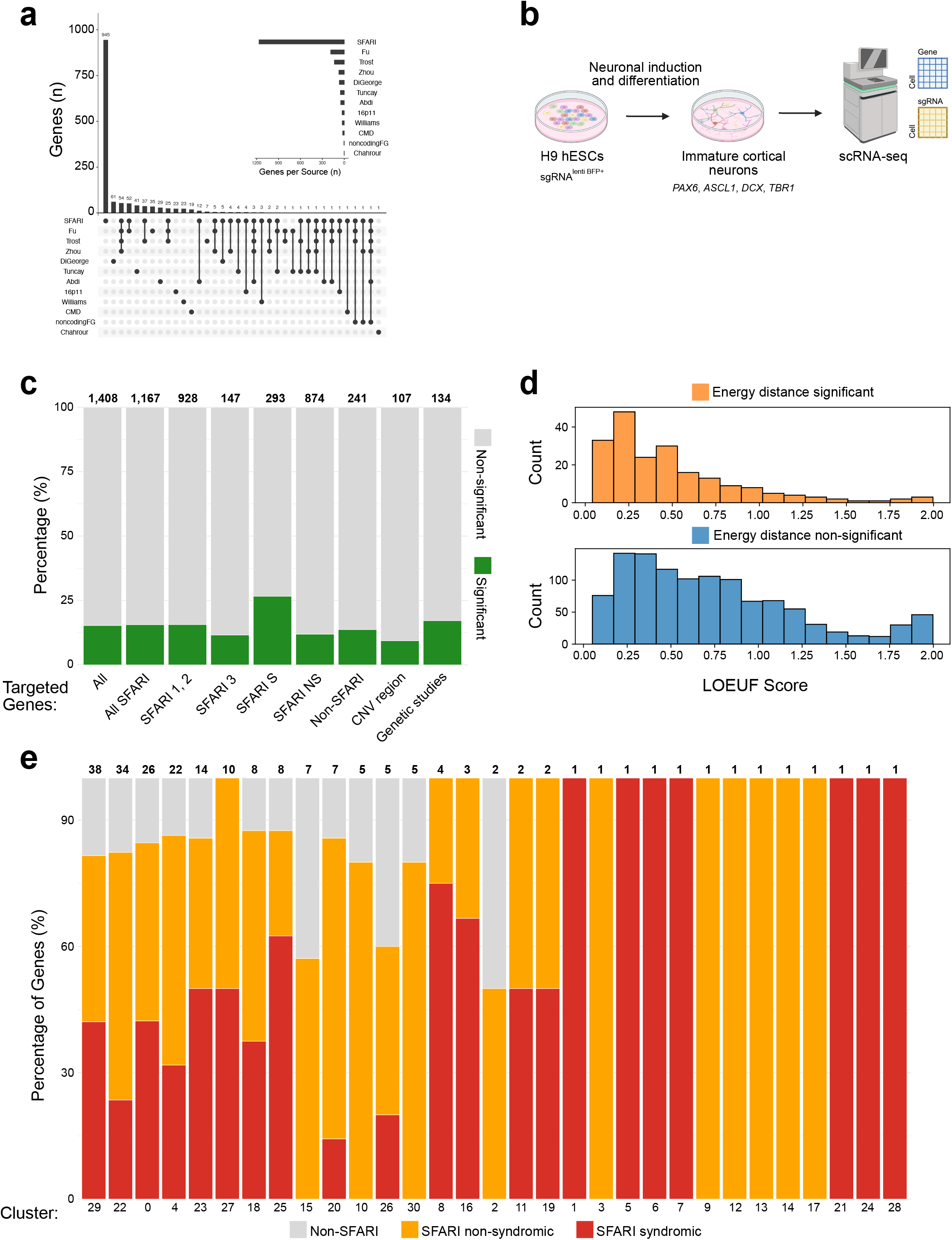
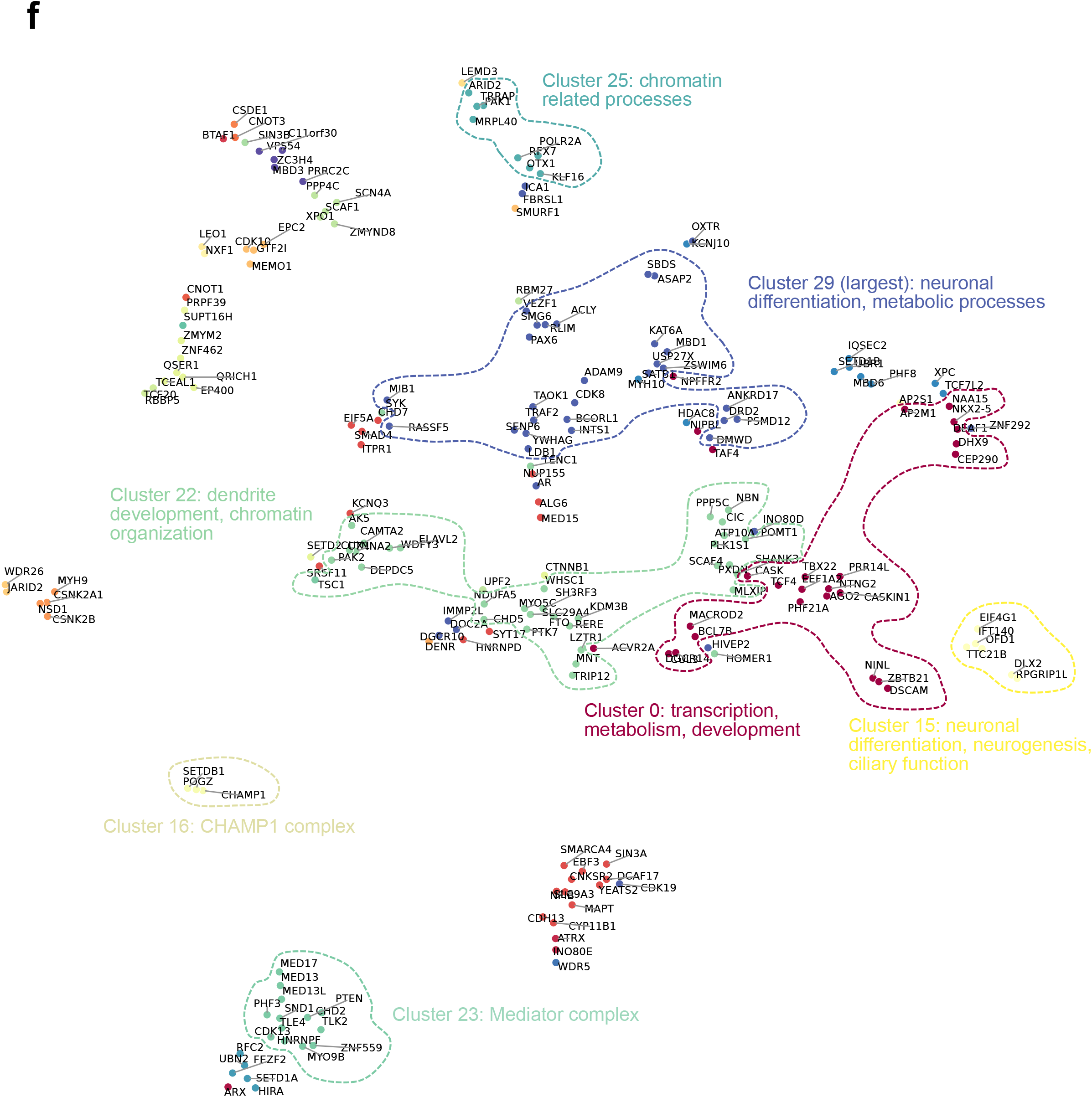
Functional characterization of ASD risk genes via CRISPRi Perturb-seq in hESC-derived cortical neurons. (a) UpSet plot illustrating the overlap among the different sources of ASD risk genes selected for perturbation. Vertical bars represent the number of genes unique to or shared by specific source combinations, indicated by connected dots below. (b) Experimental schematic of the CRISPRi Perturb-seq workflow. sgRNAs targeting ASD risk genes and non-targeting controls were transduced into hESCs, followed by differentiation into cortical neurons and subsequent scRNA-seq readout. (c) Proportions of energy distance-significant perturbations across gene categories. Bar height represents the percentage of energy distance-significant hits (green) within each category. Total gene counts for each category are indicated above each bar. (d) Distribution of LOEUF scores for energy distance-significant hits (top, orange) compared to non-significant genes (bottom, blue). Lower LOEUF scores indicate greater intolerance to loss-of-function variation. (e) Stacked barplots showing the composition of each of the 31 identified energy distance clusters. Total gene membership is indicated above each bar, with segments representing the percentage of syndromic SFARI genes (red), non-syndromic SFARI genes (orange), and non-SFARI genes (grey). (f) Dimensionality reduction embedding of energy distance-significant perturbations colored by cluster assignment, with key clusters of functional interest highlighted.

To identify perturbations that significantly alter the transcriptome relative to non-targeting control (NTC) cells, we performed energy distance analysis^37,42^. We removed 609 outlier targeting sgRNAs (6.4%) and 41 outlier non-targeting sgRNAs (7%) (Supplementary Table S3). Out of the 1,408 targeted ASD risk genes, 215 genes (15.3% overall hit rate) were identified as energy distance-significant hits (Figure 1c, Supplementary Tables S4 and S5). Among the perturbed SFARI genes, 15.6% were identified as significant, with higher hit rates among genes with SFARI scores of 1 or 2 (15.6%) than among genes with a score of 3 (11.6%). Syndromic SFARI genes showed the highest enrichment, with 26.7% classified as significant hits. Among non-SFARI perturbed genes, those curated from genetic studies had a higher hit rate (17.2%) than genes derived from CNV loci (9.3%). The distribution of loss-of-function observed/expected upper bound fraction (LOEUF) scores of perturbed genes from gnomAD^63^ differed significantly between energy distance-significant and non-significant genes (*P* < 0.001) (Figure 1d). Significant genes exhibited lower LOEUF scores overall, indicating greater intolerance to loss-of-function variation (Figure 1d).

The 215 significant genes organized into 31 functional clusters based on transcriptomic similarity (Figures 1e and 1f). Most clusters contained a mixture of syndromic SFARI, non-syndromic SFARI, and non-SFARI genes, indicating that transcriptional responses were not segregated by gene category (Figure 1e). Several clusters captured known functional relationships. Cluster 16 contained members of the CHAMP1 protein complex, including *SETDB1*, *POGZ*, and *CHAMP1*^64^, while cluster 23 included Mediator complex components *MED17*, *MED13*, and *MED13L*^65^ (Figure 1f). The largest cluster (cluster 29; 38 genes) was enriched for genes involved in neuronal differentiation and metabolic processes. Cluster 15 included three SFARI genes (*DLX2*, *OFD1*, and *EIF4G1*) together with three non-SFARI genes (*RPGRIP1L*, *TTC21B*, and *IFT140*) that shared transcriptional profiles associated with neuronal differentiation, neurogenesis, and ciliary function. *MRPL40*, located within the DiGeorge syndrome CNV locus, clustered with seven SFARI genes in cluster 25, which was enriched for chromatin-related processes. Cluster 0 (26 genes) was associated with fundamental cellular processes, including transcription, metabolism, and development, whereas cluster 22 (34 genes) was enriched for genes involved in dendrite development and chromatin organization.

### Majority of ASD risk genes reduce the rate of neuronal differentiation

After removal of outlier sgRNAs and cells lacking retained sgRNAs following energy distance analysis, the final dataset contained 549,727 cells (mean of 3,053 transcriptome UMIs, 1,247 sgRNA UMIs, and 3.8 sgRNAs per cell) and 9,496 sgRNAs and was used for all downstream analyses. We confirmed efficient on-target repression and high reproducibility across independent libraries (Supplementary Figure S2). Unsupervised clustering resolved three transcriptionally distinct populations: neural progenitor cells (NPCs; n = 167,619) defined by expression of *PAX6* and *OTX2*; an intermediate population (n = 267,702) expressing both NPC markers (*PAX6*, *OTX2*) and immature neuronal markers (*DCX*, *TUBB3*); and immature cortical neurons (n = 114,406) defined by expression of *DCX* and *TUBB3* (Figures 2a, 2b, and 2c). Cells carrying NTC sgRNAs (n = 59,714) were distributed across all three populations (n = 18,085 NPCs, 28,493 intermediate cells, and 13,136 immature neurons) (Figure 2d).

**Figure 2.**
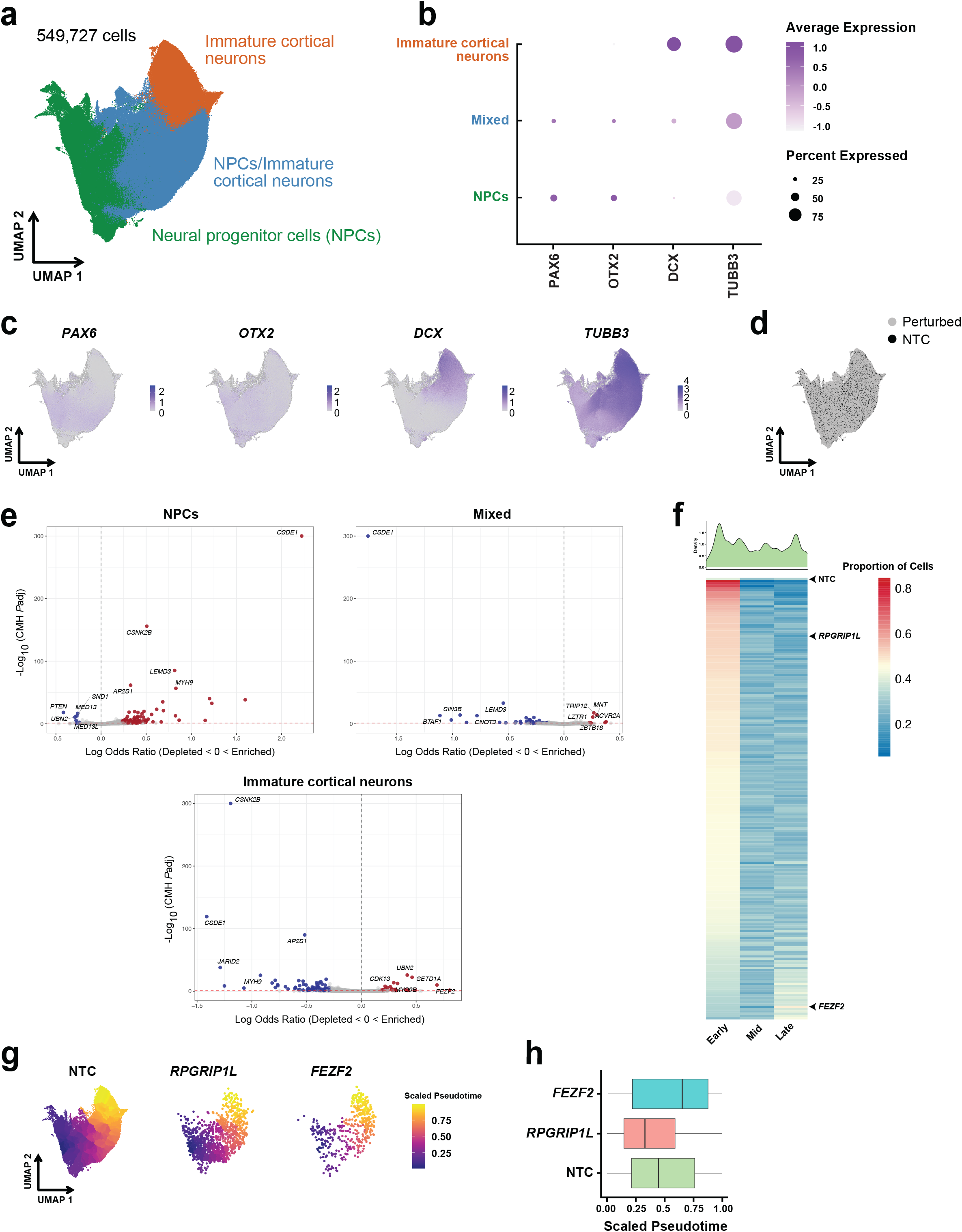
Perturbation of ASD risk genes alters cell type composition. (a) UMAP of 549,727 cells colored by cluster identity and annotated based on known cell identity markers. (b) Dot plot showing relative expression of marker genes for each cell type. Dot size represents the percentage of cells expressing each marker and dot color intensity represents average expression (light, low expression; dark, high expression). (c) UMAP showing the expression of cell identity marker genes. (d) UMAP showing the distribution of 59,714 NTC cells. (e) Volcano plots showing perturbations that significantly alter cell proportions within each cluster relative to NTC (red, enriched; blue, depleted). (f) Density distribution of NTC cells along scaled pseudotime (top). Heatmap showing the proportion of cells within early, mid, and late pseudotime bins for perturbations that significantly alter cell proportions relative to NTC (bottom). The color scale indicates the proportion of cells in each bin, normalized by row (blue, low proportion; red, high proportion). (g) UMAPs colored by pseudotime (early, blue; late, yellow) for NTC, *RPGRIP1L*, and *FEZF2* perturbations. The NPC cluster was used as the starting point for trajectory inference. (h) Median pseudotime values for NTC, *RPGRIP1L*, and *FEZF2* perturbations.

We next asked whether individual perturbations altered cell type proportions relative to NTC cells as an indicator of disrupted neuronal differentiation. By comparing the cell proportions for each perturbation with the NTC baseline, we identified 192 perturbations that produced significant compositional changes (Figure 2e, Supplementary Table S6). Of these, 149 perturbations produced cell-state distributions consistent with decreased differentiation, whereas 28 perturbations produced cell-state distributions consistent with increased differentiation. The remaining 15 perturbations were not assigned to either group because they exhibited a significant increase or decrease in the proportion of cluster 2 (intermediate state) without significant changes in the proportion of NPCs or neurons, making their effects on differentiation difficult to interpret. Several non-SFARI genes, including *INO80E*, *MRPL40*, *RPGRIP1L*, and *TTC21B*, also significantly altered cell proportions. The analysis recovered genes previously implicated in regulating neuronal differentiation, including *CSNK2A1*, *JARID2*, and *TCF7L2*^66–68^, while also identifying candidate regulators not previously linked to neuronal differentiation, including *AP2M1*, *AP2S1*, *RPGRIP1L*, *TTC21B*, and *WDR26*, whose perturbation significantly altered cell type proportions. These compositional changes were also reflected in transcriptional trajectories across pseudotime (Figure 2f). NTC cells were distributed across the full developmental trajectory (Figures 2f and 2g). In contrast, cells from perturbations associated with decreased differentiation, such as *RPGRIP1L*, had lower median pseudotime values, whereas cells from perturbations associated with increased differentiation, such as *FEZF2*, exhibited higher median pseudotime values (Figures 2f-h). Together, these findings demonstrate that perturbation of a subset of ASD risk genes significantly alters neuronal cell-state composition and pseudotime progression, revealing both established and previously unrecognized regulators of neuronal differentiation.

### Distinct ASD risk genes converge on shared neurodevelopmental gene programs

To identify coordinated transcriptional responses across perturbations, we applied consensus non-negative matrix factorization (cNMF)^46^, yielding 200 distinct gene programs (GPs) (Supplementary Figures S3a-d, Supplementary Tables S7-S9). On average, each GP was significantly dysregulated by 12.5 perturbations, while each perturbation significantly dysregulated an average of 8.5 GPs (Supplementary Figures S3e-f, Supplementary Tables S10 and S11). Of the 200 GPs, 18 were significantly enriched for SFARI ASD risk genes based on their top driver genes (Supplementary Table S11), identifying transcriptional programs that are particularly vulnerable to ASD-associated genetic disruption.

GP1 was the most frequently dysregulated program and the most commonly downregulated, altered by 132 perturbations, including 122 that downregulated and 10 that upregulated its activity (Figure 3a). Gene ontology (GO) analysis showed that GP1 is significantly enriched for microtubule polymerization and depolymerization, neuroepithelial cell differentiation, and nervous system development (Figure 3b). Its top five driver genes (*TUBB3*, *TAGLN3*, *CRABP1*, *TUBA1A*, and *STMN2*) are each essential for neuronal differentiation. GP1 expression was largely restricted to immature cortical neurons (Figure 3c). The strongest downregulators of GP1 were perturbations of *CSDE1*, *BTAF1*, *SIN3B*, *WDR26*, and *CNOT3*; the strongest upregulators were *MED13L*, *CDK13*, *MYO9B*, *UBN2*, and *TLE4* (Supplementary Figure S4a). On the other hand, the most frequently upregulated program was GP18, which was significantly enriched for SFARI ASD risk genes and associated with dopaminergic neuron differentiation, transcription, and cell-cell adhesion (Supplementary Figures S4b and S4c). GP18 was significantly regulated by 23 perturbations, with *WDR5*, *ATRX*, *INO80E*, *CYP11B1*, and *MAPT* producing the strongest upregulation in program activity. Its principal driver genes included *SLIT2*, *TTC6*, *SHH*, *FOXA1*, and *PDZRN4*.

**Figure 3.**
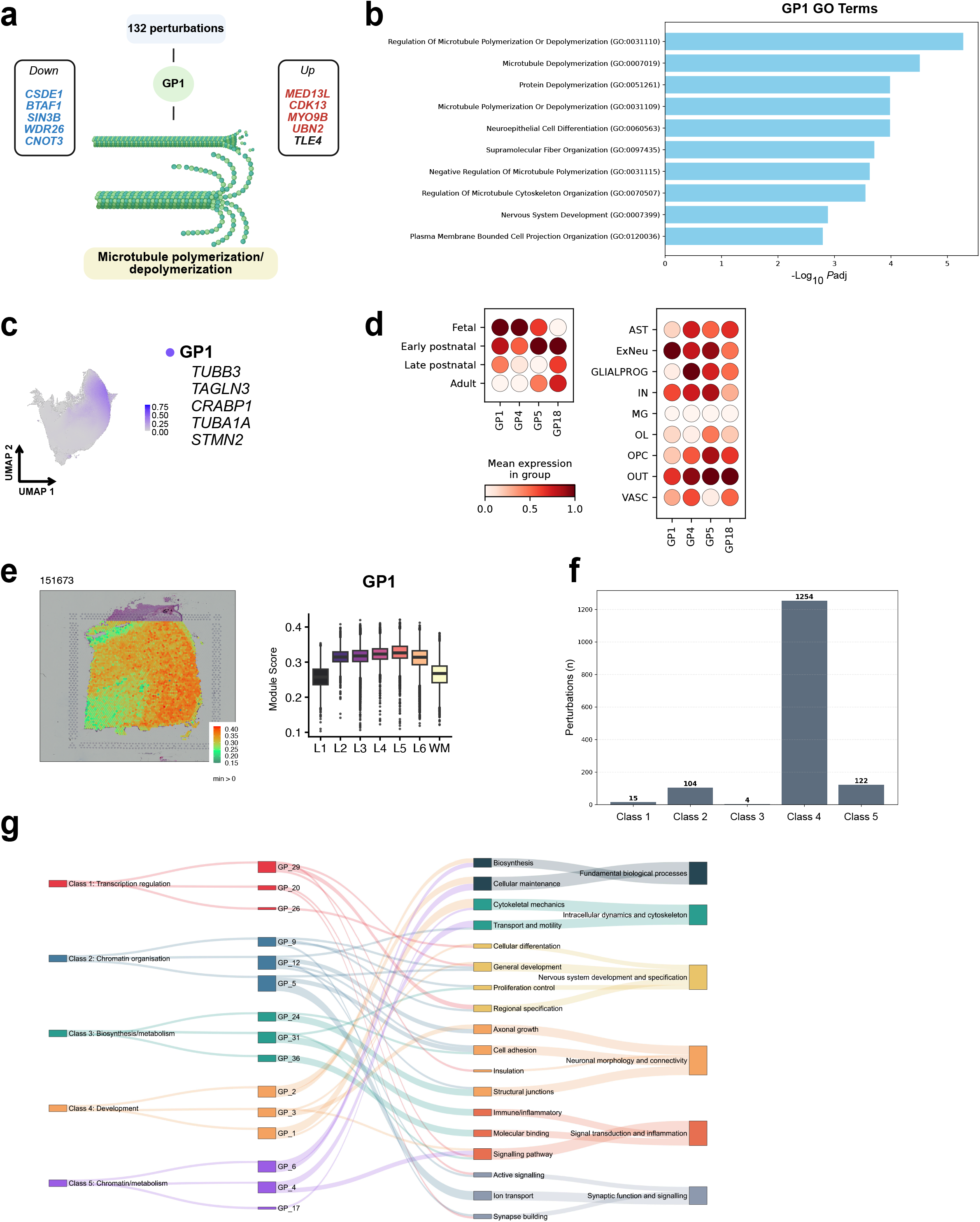
Gene programs reveal convergent biological pathways across diverse perturbations. (a) Schematic overview of GP1 showing the total number of perturbations that significantly dysregulate GP1, the top 5 down- and upregulating perturbations, and its primary biological function. (b) GO analysis showing the top 10 significantly enriched biological processes for GP1. (c) UMAP showing GP1 activity. (d) Dot plot showing relative GP activity across four developmental age ranges and across different cell types in the human cortical development dataset^48^. Dot color intensity indicates mean expression level. (e) Spatial expression map showing GP1 scores projected onto a representative tissue section (sample 151673). Spatial coordinates are aligned with the underlying tissue spanning the six cortical layers (L1-L6) and white matter (WM) (left). Boxplot showing GP1 scores across cortical layers and WM, aggregated across all spatial transcriptomics samples (right). (f) Barplot showing the distribution and total count of perturbations assigned to each mixture model class. (g) Sankey plot illustrating convergence of perturbations onto core biological pathways. Flow lines track from mixture model classes with functional annotation (left), through their top 3 driving GPs (middle left), into 18 functional subcategories (middle right), and converging into 6 major biological categories (right). The width of ribbons and vertical blocks indicate total counts of associated elements at each level.

Two additional programs GPs, GP4 and GP5, were significantly enriched for SFARI ASD risk genes and highlighted biological pathways recurrently disrupted across ASD-associated perturbations. GP4, associated with cell migration, signal transduction, and Wnt signaling, was upregulated by 58 perturbations (top: *RPGRIP1L*, *DLX2*, *WDR26*, *CSNK2B*, and *TTC21B*) and downregulated by 17 perturbations (top: *CSDE1*, *NXF1*, *RBP5*, *CHD2*, and *MEMO1*), with driver genes *IGFBP5*, *SMOC1*, *NRXN3*, *ZNF385B*, and *TSHZ2* (Supplementary Figures S4d and S4e). GP5 associated with transmembrane ion transport, nervous system development, and cell-cell adhesion, was upregulated by 8 perturbations (top: *FEZF2*, *CDK13*, *UBN2*, *MYO9B*, and *PHF3*) and downregulated by 66 perturbations (top: *CSDE1*, *SIN3B*, *BTAF1*, *WDR26*, and *CNOT3*), with driver genes *GRIA1*, *LSAMP*, *FAT3*, *ENOX2*, and *CNTN4* (Supplementary Figures S4f and S4g). These data indicate that diverse ASD risk genes converge on a relatively small number of shared transcriptional programs governing fundamental neurodevelopmental processes.

To assess the developmental relevance of these *in vitro* GPs, we projected them onto a published single-cell RNA-sequencing dataset spanning prenatal and postnatal human cortical development^48^, tracking their expression across four major developmental stages. GP usage scores revealed distinct temporal patterns across development (Figure 3d). GP1 and GP4 were most highly expressed during fetal development, while GP5 and GP18 peaked in activity during early postnatal development (Figure 3d). We also examined GP activity across cell types using snRNA-seq and spatial transcriptomics datasets from the human cortex^48,49^. GP1 showed the highest activity in excitatory neurons (Figure 3d), and spatial transcriptomics mapping localized GP1 activity across cortical layers L2 to L6 (Figure 3e). GP4, GP5, and GP18 showed activity across the majority of cortical cell types, with the exception of microglia and oligodendrocytes (Figure 3d). Together, these findings indicate that the GPs identified in our screen correspond to transcriptional programs that are active across distinct cell types and cortical layers during human cortical development *in vivo*.

Given this convergence at the level of individual GPs, we next asked whether perturbations could be grouped into higher-order functional classes based on their GP dysregulation profiles. Latent mixture modeling^69^ identified five distinct perturbation classes (Supplementary Figures S5a-f). Class sizes ranged from 4 perturbations (class 3) to 1,254 perturbations (class 4) (Figure 3f, Supplementary Table S12). GO analysis demonstrated that each class was characterized by distinct biological functions: class 1 (15 perturbations) was enriched for transcriptional regulation, class 2 (104 perturbations) for chromatin organization, class 3 (4 perturbations) for biosynthesis and metabolism, class 4 (1,254 perturbations) for development, and class 5 (122 perturbations) for chromatin- and metabolism-related processes.

To map the biological themes underlying these perturbation classes, we identified the three highest-contributing GPs within each class (Supplementary Figure S5g, Supplementary Table S13). Unlike the other classes, class 4, the largest, lacked strongly dominant GPs, exhibiting uniformly low GP z-scores and suggesting that it represents a common baseline transcriptional response across diverse genetic perturbations, rather than a specialized cellular state. Integrating perturbation classes, their dominant GPs, and functional categories (Figure 3g, Supplementary Tables S13 and S14) revealed eighteen subcategories that grouped into six overarching biological themes: fundamental biological processes, intracellular dynamics and cytoskeleton, nervous system development and specification, neuronal morphology and connectivity, signal transduction and inflammation, and synaptic function and signaling. The extensive overlap between convergent trajectories in this highly interconnected network demonstrates that regardless of an ASD risk gene’s primary function, even when distinct transcriptional programs are initially targeted, their downstream effects consistently converge on a limited set of core neurodevelopmental processes. For example, ASD risk genes *KAT6A* (chromatin regulation), *CNOT3* (metabolism), and *CDK10* (transcriptional regulation), belonging to classes 5, 3, and 1, respectively, converge on signal transduction through distinct GPs (GP4, GP31, and GP29), illustrating how perturbations with different primary functional associations can converge on a shared biological theme.

### Loss of *CHAMP1* downregulates the Wnt signaling pathway

Among the 200 GPs, we identified four (GP4, GP19, GP25, and GP40) significantly enriched for Wnt signaling-related GO terms (Figure 4a). Interestingly, each GP was defined by a distinct set of Wnt pathway-associated driver genes. GP4 included *GLI3*, *GPC3*, *LEF1*, *SFRP1*, *SFRP2*, and *SULF1*; GP19 included *COL1A1*, *CPE*, *GREM1*, and *SFRP4*; GP25 included *BMP2*, *IGFBP2*, *TPBG*, *WNT1*, *WNT10B*, *WNT3A*, and *WNT4*; and GP40 included *CDH3* and *GATA3* (Figure 4a, Supplementary Table S15). We next identified perturbations that significantly upregulate or downregulate each Wnt-associated GP. Across GP4, GP19, GP25, and GP40, 58, 44, 18, and 11 perturbations significantly upregulated GP activity, respectively. *CSNK2B* and *JARID2* were the only perturbations that consistently upregulated all four Wnt-associated GPs. In contrast, 17, 5, 2, and 4 perturbations significantly downregulated GP4, GP19, GP25, and GP40, respectively, with *CHAMP1* perturbation representing the sole shared downregulator across all four Wnt-associated GPs (Supplementary Table S15). While CSNK2B and JARID2 have previously been implicated in Wnt pathway regulation^70,71^, CHAMP1 has not. Together, these findings demonstrate a strong functional convergence on the Wnt signaling pathway, revealing that disruption of numerous ASD risk genes affects this critical pathway, and uncovering CHAMP1 as a previously unrecognized regulator of this pathway.

**Figure 4.**
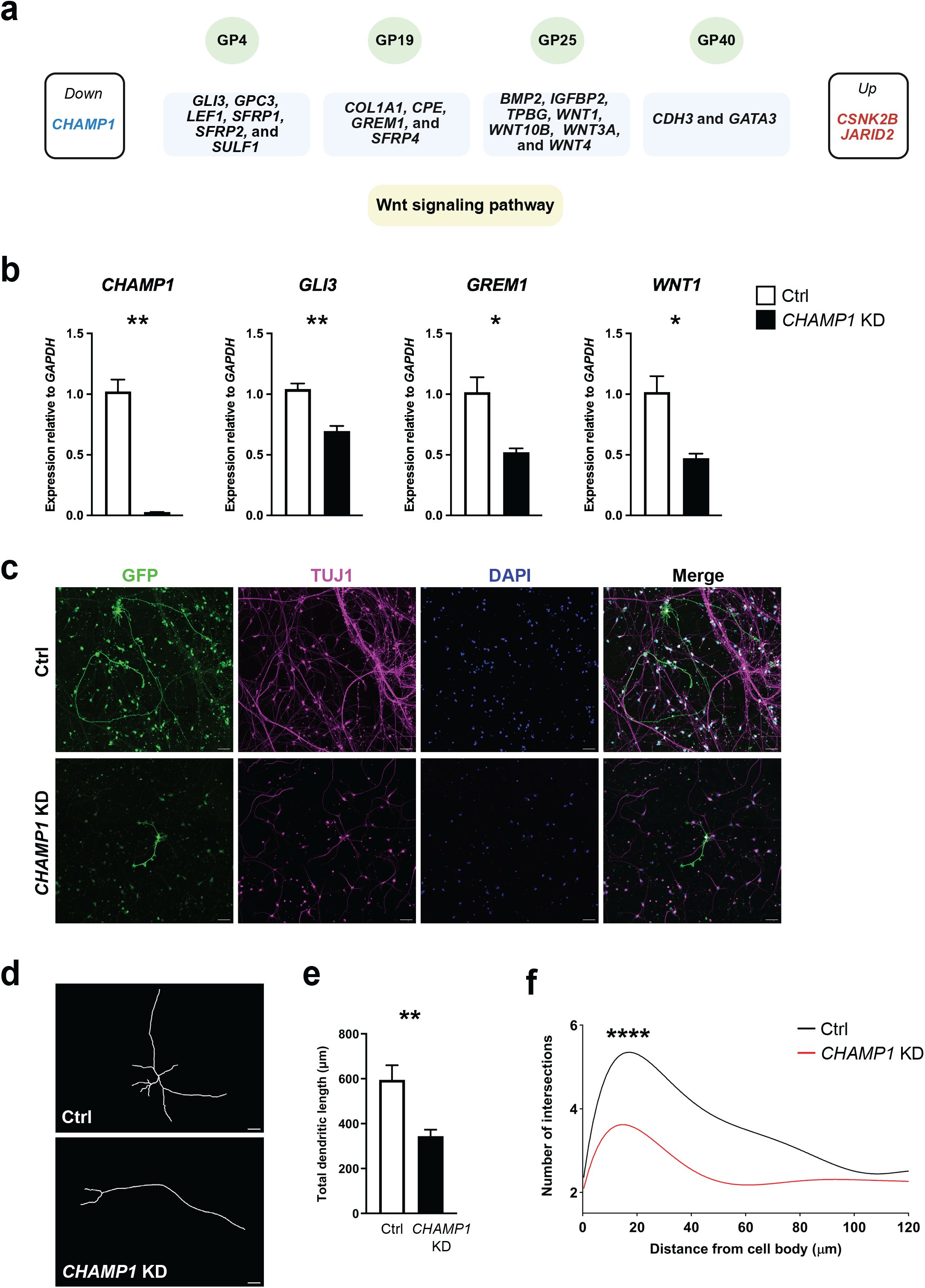
Functional convergence of ASD risk genes on the Wnt signaling pathway. (a) Schematic overview of the functional convergence of four GPs on the Wnt signaling pathway. Individual driver genes are mapped under each program panel, and adjacent panels display the one shared downregulator and two shared upregulators across all four GPs. (b) qRT-PCR validation of *CHAMP1* knockdown (KD) and downregulation of the Wnt-associated genes *GLI3* for GP4, *GREM1* for GP19, and *WNT1* for GP25 (*CHAMP1*: \*\**P* = 0.0098, *GLI3*: \*\**P* = 0.0054, *GREM1*: \**P* = 0.0173, *WNT1*: \**P* = 0.0159; n = 3 Ctrl, 3 KD). Values are mean ± SEM. Data were analyzed using two-tailed unpaired t-test (*GLI3*, *GREM1*, *WNT1*) or Welch’s t-test (*CHAMP1*). (c) Representative immunofluorescence images of Ctrl and *CHAMP1* KD neurons showing GFP, the neuronal marker TUJ1, and DAPI. Scale bars, 50 μm. (d) Representative neuronal tracings of GFP-transfected Ctrl and *CHAMP1* KD neurons. Scale bars, 20 μm. (e) Quantification of total dendritic length showing a significant decrease in *CHAMP1* KD neurons (\*\**P* = 0.0019; n = 18 Ctrl, 21 Perturbed). Values are mean ± SEM. Data were analyzed using Welch’s t-test. (f) Sholl analysis showing a significant decrease in the complexity of *CHAMP1* KD neurons (\*\*\*\**P* < 0.0001; n = 18 Ctrl, 21 Perturbed). Data were analyzed using two-way ANOVA.

To independently validate CHAMP1 as a regulator of Wnt pathway-associated gene programs, we generated a *CHAMP1* knockdown (KD) cell line and confirmed efficient repression of *CHAMP1* expression by qRT-PCR (Figure 4b). We next examined representative driver genes from three Wnt-associated GPs and found that *GLI3* (GP4), *GREM1* (GP19), and *WNT1* (GP25) were each significantly downregulated in *CHAMP1* KD cells, consistent with the Perturb-seq results (Figure 4b). Because Wnt signaling plays a critical role in neuronal differentiation^72–74^, we assessed neuronal morphology following *CHAMP1* KD. We found that CHAMP1 KD neurons exhibited significant reductions in both dendritic length and dendritic complexity compared with control neurons (Figures 4c-e). These findings validate CHAMP1 as a regulator of Wnt signaling pathway-associated transcriptional programs and suggest that disruption of *CHAMP1* impairs neuronal development through altered Wnt signaling.

### Transcriptional dysregulation of coding, non-coding, and repetitive elements across ASD risk gene perturbations

We performed differential gene expression analysis to characterize the downstream transcriptional consequences of each ASD risk gene perturbation across protein-coding genes (PCGs), long non-coding RNAs (lncRNAs), and transposable elements (TEs). Across all 1,408 perturbations, we identified 105,591 total differentially expressed genes (DEGs), with an average of 69 DEGs per perturbed gene (Supplementary Tables S16 and S17). The perturbations producing the largest transcriptional responses were *CSDE1*, *SIN3B*, *CNOT3*, *BTAF1*, and *LEMD3* (Figure 5a). DEGs from 521 perturbations were significantly enriched for SFARI ASD risk genes (Figure 5b), indicating widespread convergence onto established ASD-associated pathways.

**Figure 5.**
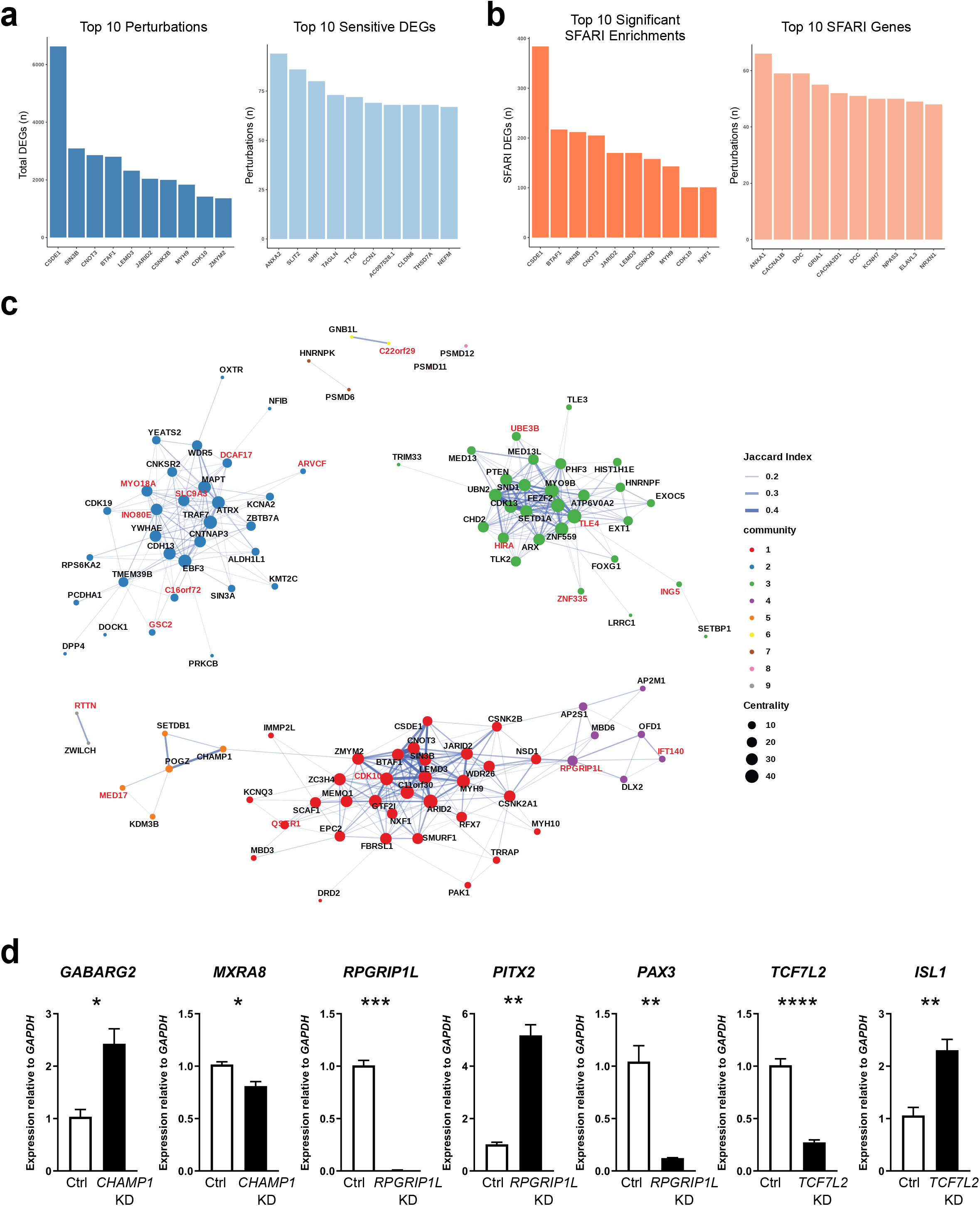
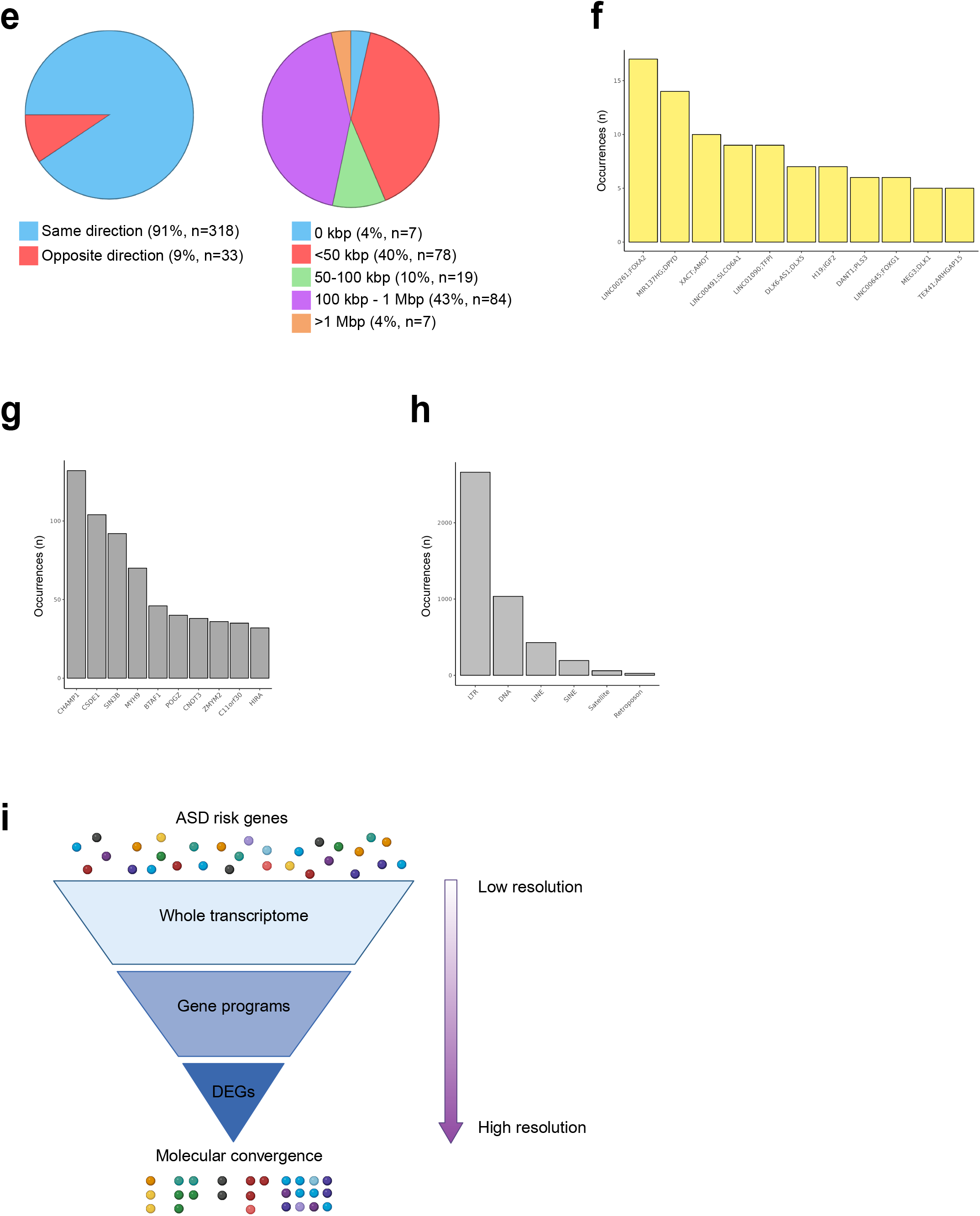
Differentially expressed genes, lncRNAs, and transposable elements across perturbations. (a) Barplots showing the top 10 perturbations with the highest number of total DEGs (left) and the top 10 most frequently dysregulated DEGs (right). (b) Barplots showing the top 10 perturbations with the highest number of DEGs that are SFARI ASD risk genes (left) and the top 10 most frequently dysregulated DEGs that are SFARI ASD risk genes (right). (c) Network plot showing directional Jaccard similarity indices between perturbation DEG lists. Only pairs with a minimum Jaccard index of 0.1 are plotted. Perturbations labeled in red are non-SFARI genes. (d) qRT-PCR validation of expression changes in downstream DEGs of *CHAMP1* (*GABARG2* \**P* = 0.0117, *MXRA8* \**P* = 0.0140), target gene repression for *RPGRIP1L* (\*\*\**P* = 0.0002) and *TCF7L2* (\*\*\*\**P* < 0.0001), and expression changes in their downstream DEGs (for *RPGRIP1L*: *PITX2* \*\**P* = 0.0015, *PAX3* \*\**P* = 0.0090; for *TCF7L2*: *ISL1*: \*\**P* = 0.0028). Values are mean ± SEM (n = 4 Ctrl, 4 KD per gene). Data were analyzed using two-tailed unpaired t-test (*GABARG2*, *MXRA8*, *TCF7L2*, *ISL1*) or Welch’s t-test (*RPGRIP1L*, *PITX2*, *PAX3*). (e) Pie charts showing the direction of dysregulation across all lncRNA-PCG co-dysregulated pairs (left) and the distribution of genomic proximity between lncRNAs and PCGs across unique co-dysregulated pairs (right). (f) Barplot showing the most frequently co-dysregulated lncRNA-PCG pairs. (g) Barplot showing the top 10 perturbations with the highest number of dysregulated transposable elements. (h) Barplot showing the frequency of dysregulated transposable element families. (i) Schematic representing the hierarchical levels of transcriptomic convergence.

Many DEGs were recurrently dysregulated across independent perturbations, revealing shared downstream transcriptional targets. The most frequently dysregulated genes were *ANXA2* (94 perturbations), *SLIT2* (86), and *SHH* (80) (Figure 5a). *ANXA2*, the most commonly dysregulated gene in the dataset, regulates cellular growth and signal transduction^75^. Its paralog, *ANXA1*, is an established ASD risk gene^76^ and was itself frequently dysregulated across 66 perturbations. Likewise, *SLIT2*, a key regulator of axon guidance and neuronal migration^77^, and *SHH*, an essential regulator of embryonic development^78^, were broadly affected across the perturbation landscape. Additional recurrently dysregulated SFARI ASD risk genes included *CACNA1B* and *DDC* (59 perturbations each), both of which play important roles in neurotransmission (Figure 5b)^79,80^. Annotation of DEG lists with established neurodevelopmental disease gene sets from BrainSpan^81,82^, lncRNAs, and transcription factors further revealed significant overlaps across multiple perturbations (Supplementary Figure S6). To quantify shared transcriptomic effects between perturbations, we performed directional Jaccard similarity analysis and identified distinct highly interconnected hubs of perturbations that shared large numbers of downstream DEGs (Figure 5c). Interestingly, several non-SFARI genes within these hubs were highly interconnected to established SFARI ASD risk genes. For example, *CDK10*, a regulator of cell growth and ciliogenesis^83^, shared many DEGs with inner nuclear membrane protein-coding *LEMD3* and chromatin regulator *ZMYM2*^84,85^. These shared transcriptional signatures through network proximity support a guilt-by-association framework, suggesting that previously uncharacterized ASD candidate risk genes converge on downstream molecular pathways disrupted by established ASD risk genes. To validate these findings, we generated *CHAMP1*, *RPGRIP1L*, and *TCF7L2* KD cell lines and measured selected DEGs by qRT-PCR. We confirmed significant changes in *GABARG2* and *MXRA8* following *CHAMP1* KD, *PITX2* and *PAX3* following *RPGRIP1L* KD, and *ISL1* following *TCF7L2* KD. In each case, the expression changes were consistent with the Perturb-seq results (Figure 5d).

We next examined coordinated dysregulation of lncRNAs and nearby PCGs. Across all perturbations, we identified 351 co-dysregulated lncRNA-PCG pairs representing 195 unique loci, defined as cases in which both the lncRNA and its nearest PCG were independently identified as DEGs within the same perturbation (Supplementary Table S18). Of these, 318 pairs showed concordant expression changes, whereas 33 showed discordant changes (Figure 5e). Genomic proximity between the lncRNAs and PCGs varied considerably, with 85 pairs within close proximity (< 50 kb), 103 pairs separated by an intermediate distance (50 kb-1 Mb), and 7 pairs separated by larger distance (> 1 Mb) (Figure 5e). Twenty of the unique PCGs are established SFARI ASD risk genes, and 12 of the unique lncRNAs overlap predicted brain-specific promoters or enhancers. We identified 4 unique lncRNA-PCG pairs (*KIF26B-AS1;HNRNPU*, *LINC00645;FOXG1*, *LINC00648;MDGA2*, and *PINCR;MAOA*) combining both features, containing an ASD risk gene adjacent to a lncRNA overlapping a predicted brain-specific enhancer. While most lncRNA-PCG pairs were observed in only a small number of perturbations, several were recurrently co-dysregulated across the dataset (Figure 5f). These included previously characterized regulatory pairs in which the lncRNA functions either in trans or as a local enhancer for the PCG: *LINC00261;FOXA1* (17 perturbations), *MIR137HG;DPYD* (14 perturbations), and *DLX6-AS1;DLX5* (7 perturbations)^86–88^. We also identified several recurrent but previously uncharacterized pairs, including *LINC00491;SLCO6A1* (9 perturbations), *DANT1;PLS3* (6 perturbations), and *LINC00645;FOXG1* (6 perturbations). Their recurrent co-dysregulation and close genomic proximity suggest that these lncRNAs may function as cis-regulatory elements during cortical development.

Finally, we examined perturbation-induced dysregulation of TEs. Across all perturbations, we identified 4,405 differentially expressed TEs (DETs), of which 4,362 were upregulated and 43 were downregulated (Supplementary Table S19). *CHAMP1* perturbation dysregulated the largest number of TEs (132 DETs) (Figure 5g). Across TE families, long terminal repeat (LTR) elements were the most frequently dysregulated TE family (2,662 DETs), followed by DNA transposons (1,035 DETs), and long interspersed nuclear elements (LINEs) (429 DETs) (Figure 5h). Together, these findings demonstrate that perturbation of ASD risk genes broadly reshapes the coding and non-coding transcriptome, revealing convergent regulation of PCGs, lncRNAs, and TEs during early cortical development.

## Discussion

We perform a large-scale CRISPRi Perturb-seq screen targeting 1,408 ASD risk genes in hESC-derived immature cortical neurons, generating a functional atlas of transcriptome-wide effects across nearly the full SFARI catalogue at single-cell resolution. Relative to prior perturbation screens in neurodevelopmental disease research^89–92^, our approach substantially expands the number of evaluated targets and improves several key design parameters. CRISPRi-mediated transcriptional repression better models haploinsufficiency than complete gene knockout, which is the predominant genetic mechanism for most ASD risk variants. Screening in a uniform cellular background at an early neurodevelopmental stage permits direct cross-perturbation comparison without the phenotypic variability inherent to organoid or assembloid systems. The single-cell transcriptomic readout captures both compositional and gene expression changes at a resolution unavailable with lower-content assays. Together, these features provide a systematic framework for comparing the consequences of genetically diverse ASD risk gene perturbations within a common human cortical neuronal context.

A persistent bottleneck in translating ASD genetic discoveries into mechanistic insights is that most catalogued risk genes identified by large-scale sequencing studies remain functionally unclassified^93^. By comprehensively profiling the entire SFARI catalogue encompassing both well-established syndromic risk genes and poorly characterized non-syndromic and candidate ASD risk genes in a single system, our dataset enables functional inference for unclassified genes based on transcriptional similarity to established disease hubs. *RPGRIP1L*, a gene involved in ciliary function^94^, illustrates this principle. Mutations in *RPGRIP1L* are primarily associated with Joubert syndrome (MIM 610937), and an inherited homozygous frameshift deletion was previously identified in an individual with ASD in our own cohort^61,95^. Here, *RPGRIP1L* clusters with established SFARI ASD risk genes *DLX2*, *OFD1*, and *EIF4G1* in cluster 15, a group involved in neuronal differentiation, neurogenesis, and ciliary processes, placing it in a defined neurodevelopmental context with known ASD risk genes. *RPGRIP1L* repression significantly increased the proportion of NPCs and decreased the proportion of neurons compared to NTC. At the GP level, *RPGRIP1L* perturbation primarily downregulated GP9 and GP5, involved in nervous system development and transmembrane ion transport, and at a single-gene resolution it dysregulated 454 downstream genes. Rather than establishing a specific molecular mechanism for *RPGRIP1L*, these convergent signatures provide testable hypotheses linking its perturbation to cortical neuronal differentiation and ciliary biology. This example illustrates how the atlas can generate mechanistic hypotheses for genes lacking functional characterization and anchor unclassified candidates within the broader disease biology.

Whether genetically heterogeneous ASD risk genes ultimately converge on shared biological pathways or drive largely independent downstream programs is a central question with direct implications for therapeutic development. Our data show that convergence operates simultaneously across multiple hierarchical levels (Figure 5i). At the broadest level, 215 of the 1,408 perturbed genes induced significant global transcriptomic dysregulation, collapsing into 31 cohesive phenotypic clusters. This finding is consistent with, and provides causal functional grounding for, the convergent co-expression and protein-protein interaction networks identified in post-mortem ASD brain studies^12–17,48^ and protein interactome screens^20^. Moving to GPs, the most broadly affected programs across perturbations (GP1, GP4, GP5) are associated with microtubule dynamics, neuronal differentiation, cell migration, and transmembrane transport, processes repeatedly implicated in ASD across independent transcriptomic and genetic studies^12–16,27^. That these programs peak during fetal and early postnatal human cortical development^48^ is consistent with the perturbations targeting developmentally relevant gene networks and processes implicated in ASD, although the present study does not establish when individual ASD risk genes exert their effects *in vivo*. Mixture modeling further organized the perturbation landscape into five perturbation classes that resolved into six major biological themes, providing a higher-order functional taxonomy for ASD risk gene biology. At single-gene resolution, DEG analysis reveals both a core of shared downstream targets and perturbation-specific signatures. Convergence is also evident at the level of cellular phenotype: of the 192 perturbations that significantly altered neuronal cell proportions, 149 were associated with decreased neuronal differentiation, indicating that disruption of diverse ASD risk genes more frequently produces cell-state distributions consistent with decreased rather than increased neuronal differentiation. Convergence and divergence coexist across the ASD risk gene landscape, operating at different levels of resolution simultaneously.

These findings fit naturally within a growing body of parallel perturbation studies in human neurodevelopmental systems^21–32^. Organoid- and assembloid-based screens have identified convergent effects on progenitor proliferation, excitatory neurogenesis, and interneuron migration^28–30^. Concurrent screens in hESC-derived cortical neurons from independent groups have recovered overlapping functional themes including chromatin regulation, RNA processing, and cytoskeletal dynamics^21–23,27^. The high-confidence ASD protein-protein interaction network map by Wang et al.^20^ centers on core processes such as cytoskeleton organization, transcriptional regulation, and RNA binding. The convergence on microtubule biology and early differentiation programs we observe is consistent with Sun et al.^27^ and Fernandez Garcia et al.^23^, reinforcing that these represent reproducible vulnerabilities in developing cortical neurons rather than platform-specific artifacts. The current screen extends these observations by evaluating 1,408 ASD risk genes within a common experimental system and by integrating transcriptional, cellular, and gene-program-level phenotypes. Our screen is distinguished by its scale, the broadest single-system evaluation of ASD risk genes to date, and by its use of CRISPRi, which models haploinsufficiency and avoids the compensatory responses that can confound complete knockouts.

Our identification of CHAMP1 as a Wnt signaling pathway regulator is among the more mechanistically tractable findings. CHAMP1 is a component of the heterochromatin assembly complex^64^ and has been linked to intellectual disability, but its role in Wnt signaling has not been described. The Wnt pathway is an established driver of cortical neuron differentiation^72–74^, and its convergent downregulation by CHAMP1, alongside upregulation by CSNK2B and JARID2, shows that ASD risk genes can modulate this pathway from multiple directions. The Wnt pathway dysregulation was recovered across four independent GPs, each defined by a distinct set of driver genes, reflecting the robustness of this finding and the pathway’s broad vulnerability to ASD-associated disruption. CHAMP1 knockdown further reduced expression of representative genes from Wnt-associated programs and altered neuronal morphology, providing orthogonal support for a role of CHAMP1 in Wnt-associated transcriptional regulation and neuronal development. The intersection with neuronal differentiation phenotypes observed across the screen, including for several genes not previously linked to this process, and the reduced dendritic length and complexity we observe following *CHAMP1* KD suggest that these transcriptional effects may functionally converge on neuronal differentiation, adding mechanistic depth to the Wnt signaling pathway finding. However, whether CHAMP1 directly regulates Wnt signaling or acts through an upstream or parallel pathway remains to be determined.

The multilevel convergence we document has a direct bearing on therapeutic strategy. At the level of GPs and shared downstream DEGs, convergence onto Wnt signaling, microtubule dynamics, or differentiation programs points toward pathway-level intervention as a testable strategy for determining whether distinct upstream genetic perturbations can be functionally rescued through shared molecular nodes. At the same time, perturbation-specific DEG signatures, present even among genes that cluster together globally, define the limits of any uniform approach. The same GPs can be dysregulated in opposite directions by different perturbations, and many perturbations dysregulate unique downstream targets within those programs. These findings argue that therapeutic strategies will likely need to balance convergent and genotype-specific mechanisms: shared pathways may provide opportunities for intervention across genetically diverse forms of ASD, whereas perturbation-specific downstream effects may require more individualized approaches.

The lncRNA and TE analyses add transcriptional complexity that most perturbation studies have not captured. Recovery of recurrent lncRNA-PCG co-dysregulated pairs, including well-characterized cis-regulatory lncRNAs such as *DLX6-AS1;DLX5* and *LINC00261;FOXA1*, validates the analytical approach. Among the novel candidates, *LINC00645;FOXG1* in particular warrants functional follow-up, given FOXG1’s established role in cortical development and neurodevelopmental disease^96^. The predominant upregulation of LTRs and DNA transposons is consistent with emerging evidence linking TE dysregulation to ASD and other neurodevelopmental conditions, though the functional consequences of this upregulation in early cortical neurons remain to be determined. Importantly, the recurrent dysregulation of non-coding elements across genetically distinct perturbations suggests that regulatory consequences of ASD risk gene disruption extend beyond PCGs and may represent an underappreciated component of convergent disease biology.

Several limitations should be noted. The screen was performed in a single hESC line, and results may not fully generalize across lines, cell types, or more mature neuronal stages. CRISPRi achieves partial repression rather than complete loss of function, which better captures haploinsufficiency but may underestimate effects of severe alleles. In addition, CRISPRi-mediated repression may not fully reproduce the molecular consequences of specific disease-associated variants, including variants that alter protein function, localization, or regulatory properties without reducing transcript abundance. Individual perturbation phenotypes, at the level of neuronal morphology, synaptic function, or *in vivo* behavior, require targeted follow-up beyond the scope of this screen. Functional interpretation of individual GPs and DEG signatures will benefit from orthogonal experimental validation. Finally, cell proportion and pseudotime analyses provide evidence for altered neuronal cell states and progression along a transcriptional trajectory but do not directly measure the kinetics of differentiation.

This study provides a multilevel functional map of ASD risk gene biology in a human cortical neuron context. By linking individual gene perturbations to cellular composition, pseudotime state, recurrent transcriptional programs, and downstream coding and non-coding targets, this dataset provides multiple levels at which genetically heterogeneous ASD risk can be functionally interpreted. The hierarchical transcriptomic convergence we document, from global transcriptome states through shared GPs to individual downstream effectors, establishes both a conceptual framework and a practical resource for identifying the shared molecular nodes most amenable to intervention, while preserving the resolution needed to address the genetic heterogeneity that defines ASD.

## Declaration of interests

The authors declare no competing interests.

## Acknowledgements

This research was supported in part by the computational resources provided by the BioHPC supercomputing facility located in the Lyda Hill Department of Bioinformatics, University of Texas Southwestern Medical Center. Some schematics in the figures were created with BioRender.com. We thank members of the Chahrour, Hon, and Munshi laboratories for critical feedback on the manuscript, and members of the IGVF Consortium Neuroscience Focus Group for insightful discussions.

## Funding statement

This work is supported by the National Institutes of Health UM1HG011996 (G.H.C., N.V.M., W.L.K., M.H.C) and the Walter and Lillian Cantor Foundation (M.H.C).

## Author contributions

M.H.C., G.C.H., N.V.M., and W.L.K. conceived the study, acquired funds, and oversaw the project. A.G., W.C.C., M.N., C.T., H.Z., L.W., S.S., M.D., and K.K. designed and performed experiments and analyzed data. A.G., W.C.C, and M.H.C. wrote the manuscript. All authors participated in reviewing and editing of the manuscript.

## Data availability

The raw and analyzed data are available through the IGVF Consortium Data Portal (https://data.igvf.org/) under accession number IGVFDS9825FOBP. Any additional information required to reanalyze the data reported in this paper is available from the corresponding author upon request.

## Code availability

The code used for data analysis in this study is available on the Chahrour lab GitHub repository at https://github.com/chahrourlab/ASD_CRISPRi_PerturbSeq/.

## Supplementary Figure Legends

**Figure S1. Quality control of neuronal differentiation.**

(a) qRT-PCR analysis of marker gene expression. The expression of the stem cell marker *OCT4* significantly decreased (\*\*\**P* = 0.0004; n = 3 Ctrl, 3 Perturbed), whereas the expression of the neuronal markers *PAX6*, *ASCL1*, *DCX*, and *TBR1* significantly increased (\*\*\*\**P* < 0.0001; n = 6 Ctrl, 6 Perturbed) after 6 days of differentiation. Values are mean ± SEM. Data were analyzed using two-tailed unpaired t-test (*PAX6*) or Welch’s t-test (*OCT4*, *ASCL1*, *DCX*, *TBR1*). (b) Immunofluorescence staining. Representative images show that OCT4 (red) was detected only in undifferentiated cells, whereas DCX (green) was detected only in differentiated cells. Scale bars, 50 μm.

**Figure S2. Quality of Perturb-seq data.**

(a) On-target repression efficiency. (b) Feature plots illustrating the distribution of cells across the 24 Perturb-seq libraries.

**Figure S3. Optimization of the number of gene programs.**

(a) Stability and error profile as a function of the number of components (K). K values tested: 50, 100, 200, 250, and 500. (b) Total number of unique GO terms recovered across all gene programs (GPs) at each tested K value. (c) Total number of unique perturbations recovered across all GPs at each tested K value. (d) Cosine similarity matrix calculated between the top 300 driver genes of each GP. (e) Histogram showing the distribution of the number of perturbations significantly dysregulating each GP at K = 200. (f) Histogram showing the distribution of the number of GPs significantly dysregulated by individual perturbations at K = 200.

**Figure S4. Functional annotation and dysregulation of gene programs.**

(a) Waterfall plot showing perturbations that significantly dysregulate GP1. The y-axis represents log_2_ fold change. Perturbations annotated as non-SFARI genes are denoted in red. (b) Waterfall plot showing perturbations that significantly dysregulate GP18. The y-axis represents log_2_ fold change. Perturbations annotated as non-SFARI genes are denoted in red. (c) GO analysis showing the top 10 significantly enriched biological processes for GP18. (d) Waterfall plot showing perturbations that significantly dysregulate GP4. The y-axis represents log_2_ fold change. Perturbations annotated as non-SFARI genes are denoted in red. (e) GO analysis showing the top 10 significantly enriched biological processes for GP4. (f) Waterfall plot showing perturbations that significantly dysregulate GP5. The y-axis represents log_2_ fold change. Perturbations annotated as non-SFARI genes are denoted in red. (g) GO analysis showing the top 10 significantly enriched biological processes for GP5.

**Figure S5. Model fit metrics for mixture model class selection.**

(a) Bayesian information criterion (BIC) curve. (b) Akaike information criterion (AIC) curve. (c) Log-likelihood curve evaluating overall model fit. (d) Model entropy curve. (e) Sample-size adjusted BIC (SABIC) curve. (f) Consistent AIC (CAIC) curve. (g) Heatmap showing the gene programs (GPs) defining the identified perturbation classes. Rows represent the five classes and columns represent the 200 GPs. Color scale indicates the average GP z-score per class, with the color intensity capped at ± 2 for visualization.

**Figure S6. Differentially expressed neurodevelopmental disease genes, transcription factors, and long non-coding RNAs.**

(a) Barplots showing the top 10 perturbations with the highest number of neurodevelopmental disorder (NDD) DEGs (left) and the top 10 most frequently dysregulated NDD DEGs (right). (b) Barplots showing the top 10 perturbations with the highest number of transcription factor (TF) DEGs (left) and the top 10 most frequently dysregulated TF DEGs (right). (c) Barplots showing the top 10 perturbations with the highest number of long non-coding RNA (lncRNA) DEGs (left) and the top 10 most frequently dysregulated lncRNA DEGs (right).

## Supplementary Table Legends

**Table S1. Summary of sgRNAs used in this study.**

**Table S2. Summary of single-cell library statistics.**

**Table S3. sgRNA outlier analysis.**

**Table S4. Energy distance permutation analysis for sgRNAs.**

**Table S5. Energy distance-significant hits.**

**Table S6. Summary of perturbations causing significant changes in cell proportions relative to NTC.**

**Table S7. Top 300 driver genes per gene program.**

**Table S8. Number of top genes per gene program determined by the knee cutoff method.**

**Table S9. Significant gene ontology terms per gene program.**

**Table S10. Gene programs significantly dysregulated by at least one perturbation.**

**Table S11. Summary of gene programs.**

**Table S12. Class assignment per perturbation using mixture modeling.**

**Table S13. Average gene program z-score per class.**

**Table S14. Gene ontology terms of the top 3 driver gene programs per class, grouped into functional categories.**

**Table S15. Summary of Wnt signaling pathway-associated gene programs.**

**Table S16. Differentially expressed genes per perturbation.**

**Table S17. Summary of differentially expressed genes per perturbation.**

**Table S18. Co-dysregulated lncRNA-PCG pairs.**

**Table S19. Differentially expressed transposable elements per perturbation.**

## Notes

### Competing Interest Statement

The authors have declared no competing interest.

## References

1 Dias, C. M. & Walsh, C. A. Recent advances in understanding the genetic architecture of autism. Annu Rev Genomics Hum Genet 21, 289–304 (2020). 10.1146/annurev-genom-121219-082309

2 Park, H. R. et al. A short review on the current understanding of autism spectrum disorders. Experimental neurobiology 25, 1–13 (2016). 10.5607/en.2016.25.1.1

3 Zeidan, J. et al. Global prevalence of autism: A systematic review update. Autism research : official journal of the International Society for Autism Research 15, 778–790 (2022). 10.1002/aur.2696

4 Bai, D. et al. Association of genetic and environmental factors with autism in a 5-country cohort. JAMA psychiatry 76, 1035–1043 (2019). 10.1001/jamapsychiatry.2019.1411

5 Cirnigliaro, M. et al. The contributions of rare inherited and polygenic risk to asd in multiplex families. Proceedings of the National Academy of Sciences of the United States of America 120, e2215632120 (2023). 10.1073/pnas.2215632120

6 Abdi, M. et al. Genomic architecture of autism spectrum disorder in qatar: The baraka-qatar study. Genome medicine 15, 81 (2023). 10.1186/s13073-023-01228-w

7 Fu, J. M. et al. Rare coding variation provides insight into the genetic architecture and phenotypic context of autism. Nature genetics 54, 1320–1331 (2022). 10.1038/s41588-022-01104-0

8 Trost, B. et al. Genomic architecture of autism from comprehensive whole-genome sequence annotation. Cell 185, 4409–4427.e4418 (2022). 10.1016/j.cell.2022.10.009

9 Zhou, X. et al. Integrating de novo and inherited variants in 42,607 autism cases identifies mutations in new moderate-risk genes. Nature genetics 54, 1305–1319 (2022). 10.1038/s41588-022-01148-2

10 Satterstrom, F. K. et al. Large-scale exome sequencing study implicates both developmental and functional changes in the neurobiology of autism. Cell 180, 568–584.e523 (2020). 10.1016/j.cell.2019.12.036

11 Abrahams, B. S. et al. Sfari gene 2.0: A community-driven knowledgebase for the autism spectrum disorders (asds). Mol Autism 4, 36 (2013). 10.1186/2040-2392-4-36

12 Gandal, M. J. et al. Broad transcriptomic dysregulation occurs across the cerebral cortex in asd. Nature 611, 532–539 (2022). 10.1038/s41586-022-05377-7

13 Parikshak, N. N. et al. Integrative functional genomic analyses implicate specific molecular pathways and circuits in autism. Cell 155, 1008–1021 (2013). 10.1016/j.cell.2013.10.031

14 Parikshak, N. N. et al. Genome-wide changes in lncrna, splicing, and regional gene expression patterns in autism. Nature 540, 423–427 (2016). 10.1038/nature20612

15 Wu, Y. E., Parikshak, N. N., Belgard, T. G. & Geschwind, D. H. Genome-wide, integrative analysis implicates microrna dysregulation in autism spectrum disorder. Nature neuroscience 19, 1463–1476 (2016). 10.1038/nn.4373

16 Wamsley, B. et al. Molecular cascades and cell type-specific signatures in asd revealed by single-cell genomics. Science (New York, NY) 384, eadh2602 (2024). 10.1126/science.adh2602

17 Voineagu, I. et al. Transcriptomic analysis of autistic brain reveals convergent molecular pathology. Nature 474, 380–384 (2011). 10.1038/nature10110

18 Pintacuda, G. et al. Protein interaction studies in human induced neurons indicate convergent biology underlying autism spectrum disorders. Cell genomics 3, 100250 (2023). 10.1016/j.xgen.2022.100250

19 Liao, C. et al. Convergent coexpression of autism-associated genes suggests some novel risk genes may not be detectable in large-scale genetic studies. Cell genomics 3, 100277 (2023). 10.1016/j.xgen.2023.100277

20 Wang, B. et al. Autism mutations rewire protein interaction networks to drive neurodevelopmental pathology. Science (New York, NY) 393, eady4523 (2026). 10.1126/science.ady4523

21 Amelan, A. et al. Crispr knockout screens reveal genes and pathways essential for neuronal differentiation and implicate peds1 in neurodevelopment. Nature neuroscience 29, 592–603 (2026). 10.1038/s41593-025-02165-0

22 Gordon, A. et al. Developmental convergence and divergence in human stem cell models of autism. Nature 651, 707–719 (2026). 10.1038/s41586-025-10047-5

23 Fernandez Garcia, M., et al. Transcriptomic and phenotypic convergence of neurodevelopmental disorder risk genes in vitro and in vivo. Nature neuroscience 29, 1079–1094 (2026). 10.1038/s41593-026-02247-7

24 Ding, J. W. et al. Dissecting gene regulatory networks governing human cortical cell fate. Nature 651, 732–742 (2026). 10.1038/s41586-025-09997-7

25 Teter, O. M. et al. Crispri-based screen of autism spectrum disorder risk genes in microglia uncovers roles of adnp in microglia endocytosis and synaptic pruning. Molecular Psychiatry 30, 4176–4193 (2025). 10.1038/s41380-025-02997-z

26 Andersen, R. E. et al. Transcriptomic convergence and the female protective effect in autism. bioRxiv : the preprint server for biology (2025). 10.1101/2025.01.20.634000

27 Sun, N. et al. Autism genes converge on microtubule biology and rna-binding proteins during excitatory neurogenesis. bioRxiv : the preprint server for biology (2024). 10.1101/2023.12.22.573108

28 Meng, X. et al. Assembloid crispr screens reveal impact of disease genes in human neurodevelopment. Nature 622, 359–366 (2023). 10.1038/s41586-023-06564-w

29 Li, C. et al. Single-cell brain organoid screening identifies developmental defects in autism. Nature 621, 373–380 (2023). 10.1038/s41586-023-06473-y

30 Paulsen, B. et al. Autism genes converge on asynchronous development of shared neuron classes. Nature 602, 268–273 (2022). 10.1038/s41586-021-04358-6

31 Jin, X. et al. In vivo perturb-seq reveals neuronal and glial abnormalities associated with autism risk genes. Science (New York, NY) 370 (2020). 10.1126/science.aaz6063

32 Cederquist, G. Y. et al. A multiplex human pluripotent stem cell platform defines molecular and functional subclasses of autism-related genes. Cell stem cell 27, 35–49.e36 (2020). 10.1016/j.stem.2020.06.004

33 Schwarz, L. A. et al. Cortical development dynamics across autism spectrum disorder mouse models. Nature 656, 159–172 (2026). 10.1038/s41586-026-10679-1

34 Stirtz, E. et al. A genetic screen reveals dosage-sensitive effects of ASD genes and identifies domino as a regulator of synaptic and behavioral phenotypes. eLife 15 (2026). 10.7554/eLife.111906.1

35 Sivakumar, S. et al. Benchmarking and optimizing perturb-seq in differentiating human pluripotent stem cells. Stem cell reports 20, 102713 (2025). 10.1016/j.stemcr.2025.102713

36 Qi, Y. et al. Combined small-molecule inhibition accelerates the derivation of functional cortical neurons from human pluripotent stem cells. Nature biotechnology 35, 154–163 (2017). 10.1038/nbt.3777

37 Replogle, J. M. et al. Mapping information-rich genotype-phenotype landscapes with genome-scale perturb-seq. Cell 185, 2559–2575.e2528 (2022). 10.1016/j.cell.2022.05.013

38 McKenna, A. & Shendure, J. Flashfry: A fast and flexible tool for large-scale crispr target design. BMC biology 16, 74 (2018). 10.1186/s12915-018-0545-0

39 Stoeckius, M. et al. Cell hashing with barcoded antibodies enables multiplexing and doublet detection for single cell genomics. Genome biology 19, 224 (2018). 10.1186/s13059-018-1603-1

40 Duan, J. & Hon, G. C. Fba: Feature barcoding analysis for single cell rna-seq. Bioinformatics (Oxford, England) 37, 4266–4268 (2021). 10.1093/bioinformatics/btab375

41 Wang, Y., Xie, S., Armendariz, D. & Hon, G. C. Computational identification of clonal cells in single-cell crispr screens. BMC genomics 23, 135 (2022). 10.1186/s12864-022-08359-1

42 Takeuchi, C. et al. Comprehensive perturbation of transcription factors in human cardiomyocytes reveals the regulatory architecture of congenital heart disease. bioRxiv : the preprint server for biology (2026). 10.64898/2025.12.15.694070

43 Takeuchi, C. et al. Regulatory network hubs guide dynamic human lineage specification. bioRxiv, 2025.2012.2015.694070 (2026). 10.64898/2025.12.15.694070

44 Cao, J. et al. The single-cell transcriptional landscape of mammalian organogenesis. Nature 566, 496–502 (2019). 10.1038/s41586-019-0969-x

45 Qiu, X. et al. Reversed graph embedding resolves complex single-cell trajectories. Nat Methods 14, 979–982 (2017). 10.1038/nmeth.4402

46 Kotliar, D. et al. Identifying gene expression programs of cell-type identity and cellular activity with single-cell rna-seq. eLife 8 (2019). 10.7554/eLife.43803

47 Satopaa, V. A., Albrecht, J. R., Irwin, D. E. & Raghavan, B. Finding a “kneedle” in a haystack: Detecting knee points in system behavior. 2011 31st International Conference on Distributed Computing Systems Workshops, 166–171 (2011).

48 Velmeshev, D. et al. Single-cell analysis of prenatal and postnatal human cortical development. Science 382, eadf0834 (2023). 10.1126/science.adf0834

49 Maynard, K. R. et al. Transcriptome-scale spatial gene expression in the human dorsolateral prefrontal cortex. Nature neuroscience 24, 425–436 (2021). 10.1038/s41593-020-00787-0

50 Tirosh, I. et al. Dissecting the multicellular ecosystem of metastatic melanoma by single-cell rna-seq. Science (New York, NY) 352, 189–196 (2016). 10.1126/science.aad0501

51 Andreatta, M. & Carmona, S. J. Ucell: Robust and scalable single-cell gene signature scoring. Computational and structural biotechnology journal 19, 3796–3798 (2021). 10.1016/j.csbj.2021.06.043

52 Morin, S. et al. Stepmix: A python package for pseudo-likelihood estimation of generalized mixture models with external variables. Journal of Statistical Software 113, 1–39 (2025). 10.18637/jss.v113.i08

53 Wang, Y. et al. Enhancer regulatory networks globally connect non-coding breast cancer loci to cancer genes. Genome biology 26, 10 (2025). 10.1186/s13059-025-03474-0

54 Drost, H. G. & Paszkowski, J. Biomartr: Genomic data retrieval with r. Bioinformatics (Oxford, England) 33, 1216–1217 (2017). 10.1093/bioinformatics/btw821

55 Ernst, J. et al. Mapping and analysis of chromatin state dynamics in nine human cell types. Nature 473, 43–49 (2011). 10.1038/nature09906

56 Cheung, I. et al. Developmental regulation and individual differences of neuronal h3k4me3 epigenomes in the prefrontal cortex. Proceedings of the National Academy of Sciences of the United States of America 107, 8824–8829 (2010). 10.1073/pnas.1001702107

57 Markenscoff-Papadimitriou, E. et al. A chromatin accessibility atlas of the developing human telencephalon. Cell 182, 754–769.e718 (2020). 10.1016/j.cell.2020.06.002

58 He, J. et al. Identifying transposable element expression dynamics and heterogeneity during development at the single-cell level with a processing pipeline scte. Nature communications 12, 1456 (2021). 10.1038/s41467-021-21808-x

59 Storer, J., Hubley, R., Rosen, J., Wheeler, T. J. & Smit, A. F. The dfam community resource of transposable element families, sequence models, and genome annotations. Mobile DNA 12, 2 (2021). 10.1186/s13100-020-00230-y

60 Tuncay, I. O. et al. The genetics of autism spectrum disorder in an east african familial cohort. Cell genomics 3, 100322 (2023). 10.1016/j.xgen.2023.100322

61 Tuncay, I. O. et al. Analysis of recent shared ancestry in a familial cohort identifies coding and noncoding autism spectrum disorder variants. NPJ genomic medicine 7, 13 (2022). 10.1038/s41525-022-00284-2

62 Vashisth, S. et al. The e3 ubiquitin ligase ube3b regulates synaptic development and cortical network activity. Autism research : official journal of the International Society for Autism Research, e70229 (2026). 10.1002/aur.70229

63 Chen, S. et al. A genomic mutational constraint map using variation in 76,156 human genomes. Nature 625, 92–100 (2024). 10.1038/s41586-023-06045-0

64 Li, F. et al. Champ1 complex directs heterochromatin assembly and promotes homology-directed DNA repair. Nature communications 16, 1714 (2025). 10.1038/s41467-025-56834-6

65 Richter, W. F., Nayak, S., Iwasa, J. & Taatjes, D. J. The mediator complex as a master regulator of transcription by rna polymerase ii. Nature reviews Molecular cell biology 23, 732–749 (2022). 10.1038/s41580-022-00498-3

66 Zhang, J., Roberts, J. M., Chang, F., Schwakopf, J. & Vetter, M. L. Jarid2 promotes temporal progression of retinal progenitors via repression of foxp1. Cell reports 42, 112237 (2023). 10.1016/j.celrep.2023.112237

67 Zhang, S. et al. Casein kinase 2 maintains the self-renewal of neural stem cells via prospero phosphorylation in drosophila. Cell & bioscience 16, 23 (2026). 10.1186/s13578-026-01532-z

68 Chodelkova, O., Masek, J., Korinek, V., Kozmik, Z. & Machon, O. Tcf7l2 is essential for neurogenesis in the developing mouse neocortex. Neural development 13, 8 (2018). 10.1186/s13064-018-0107-8

69 Litman, A. et al. Decomposition of phenotypic heterogeneity in autism reveals underlying genetic programs. Nature genetics 57, 1611–1619 (2025). 10.1038/s41588-025-02224-z

70 Asif, M. et al. De novo variants of csnk2b cause a new intellectual disability-craniodigital syndrome by disrupting the canonical wnt signaling pathway. HGG advances 3, 100111 (2022). 10.1016/j.xhgg.2022.100111

71 Adhikari, A. & Davie, J. Jarid2 and the prc2 complex regulate skeletal muscle differentiation through regulation of canonical wnt signaling. Epigenetics & chromatin 11, 46 (2018). 10.1186/s13072-018-0217-x

72 Inestrosa, N. C. & Varela-Nallar, L. Wnt signalling in neuronal differentiation and development. Cell and tissue research 359, 215–223 (2015). 10.1007/s00441-014-1996-4

73 Kim, H. et al. Dual function of wnt signaling during neuronal differentiation of mouse embryonic stem cells. Stem cells international 2015, 459301 (2015). 10.1155/2015/459301

74 Munji, R. N., Choe, Y., Li, G., Siegenthaler, J. A. & Pleasure, S. J. Wnt signaling regulates neuronal differentiation of cortical intermediate progenitors. The Journal of neuroscience : the official journal of the Society for Neuroscience 31, 1676–1687 (2011). 10.1523/jneurosci.5404-10.2011

75 White, Z. B., 2nd, Nair, S. & Bredel, M. The role of annexins in central nervous system development and disease. Journal of molecular medicine (Berlin, Germany) 102, 751–760 (2024). 10.1007/s00109-024-02443-7

76 Correia, C. T. et al. Recurrent duplications of the annexin a1 gene (anxa1) in autism spectrum disorders. Molecular autism 5, 28 (2014). 10.1186/2040-2392-5-28

77 Sherchan, P., Travis, Z. D., Tang, J. & Zhang, J. H. The potential of slit2 as a therapeutic target for central nervous system disorders. Expert opinion on therapeutic targets 24, 805–818 (2020). 10.1080/14728222.2020.1766445

78 Charytoniuk, D. et al. Sonic hedgehog signalling in the developing and adult brain. Journal of physiology, Paris 96, 9–16 (2002). 10.1016/s0928-4257(01)00075-4

79 Bunda, A. et al. Cacna1b alternative splicing impacts excitatory neurotransmission and is linked to behavioral responses to aversive stimuli. Molecular brain 12, 81 (2019). 10.1186/s13041-019-0500-1

80 Lotsios, N. S. et al. Expression of human l-dopa decarboxylase (ddc) under conditions of oxidative stress. Current issues in molecular biology 45, 10179–10192 (2023). 10.3390/cimb45120635

81 Hawrylycz, M. J. et al. An anatomically comprehensive atlas of the adult human brain transcriptome. Nature 489, 391–399 (2012). 10.1038/nature11405

82 Miller, J. A. et al. Transcriptional landscape of the prenatal human brain. Nature 508, 199–206 (2014). 10.1038/nature13185

83 Guen, V. J. et al. A homozygous deleterious cdk10 mutation in a patient with agenesis of corpus callosum, retinopathy, and deafness. American journal of medical genetics Part A 176, 92–98 (2018). 10.1002/ajmg.a.38506

84 Li, W. et al. The inner nuclear membrane protein lemd3 organizes the 3d chromatin architecture to maintain vascular smooth muscle cell identity. Nature communications 16, 8826 (2025). 10.1038/s41467-025-63876-3

85 Lezmi, E. et al. The chromatin regulator zmym2 restricts human pluripotent stem cell growth and is essential for teratoma formation. Stem cell reports 15, 1275–1286 (2020). 10.1016/j.stemcr.2020.05.014

86 Dhamija, S. et al. Linc00261 and the adjacent gene foxa2 are epithelial markers and are suppressed during lung cancer tumorigenesis and progression. Non-coding RNA 5 (2018). 10.3390/ncrna5010002

87 Feng, N. et al. Schizophrenia risk-associated snps affect expression of microrna 137 host gene: A postmortem study. Human molecular genetics 33, 1939–1947 (2024). 10.1093/hmg/ddae130

88 Ghafouri-Fard, S. et al. Dlx6-as1: A long non-coding rna with oncogenic features. Frontiers in cell and developmental biology 10, 746443 (2022). 10.3389/fcell.2022.746443

89 Ahmed, M., Muffat, J. & Li, Y. Understanding neural development and diseases using crispr screens in human pluripotent stem cell-derived cultures. Frontiers in cell and developmental biology 11, 1158373 (2023). 10.3389/fcell.2023.1158373

90 Zheng, X., Li, J. & Jin, X. Functional neurogenomics to dissect disease mechanisms across models. Annual review of genomics and human genetics 26, 189–216 (2025). 10.1146/annurev-genom-120823-125811

91 Damianidou, E., Mouratidou, L. & Kyrousi, C. Research models of neurodevelopmental disorders: The right model in the right place. Frontiers in neuroscience 16, 1031075 (2022). 10.3389/fnins.2022.1031075

92 Li, K., Ouyang, M., Zhan, J. & Tian, R. Crispr-based functional genomics screening in human-pluripotent-stem-cell-derived cell types. Cell genomics 3, 100300 (2023). 10.1016/j.xgen.2023.100300

93 Kim, S. W. & An, J. Y. Advancing precision diagnosis in autism: Insights from large-scale genomic studies. Molecules and cells 48, 100248 (2025). 10.1016/j.mocell.2025.100248

94 Wiegering, A., Rüther, U. & Gerhardt, C. The ciliary protein rpgrip1l in development and disease. Developmental biology 442, 60–68 (2018). 10.1016/j.ydbio.2018.07.024

95 Brancati, F. et al. Rpgrip1l mutations are mainly associated with the cerebello-renal phenotype of joubert syndrome-related disorders. Clinical genetics 74, 164–170 (2008). 10.1111/j.1399-0004.2008.01047.x

96 Hettige, N. C. & Ernst, C. Foxg1 dose in brain development. Frontiers in pediatrics 7, 482 (2019). 10.3389/fped.2019.00482

