## Supplementary figures and images for "Systematic CRISPRi perturbation of 1,408 autism risk genes maps multilevel transcriptional convergence in human cortical neurons"

### Figure S1

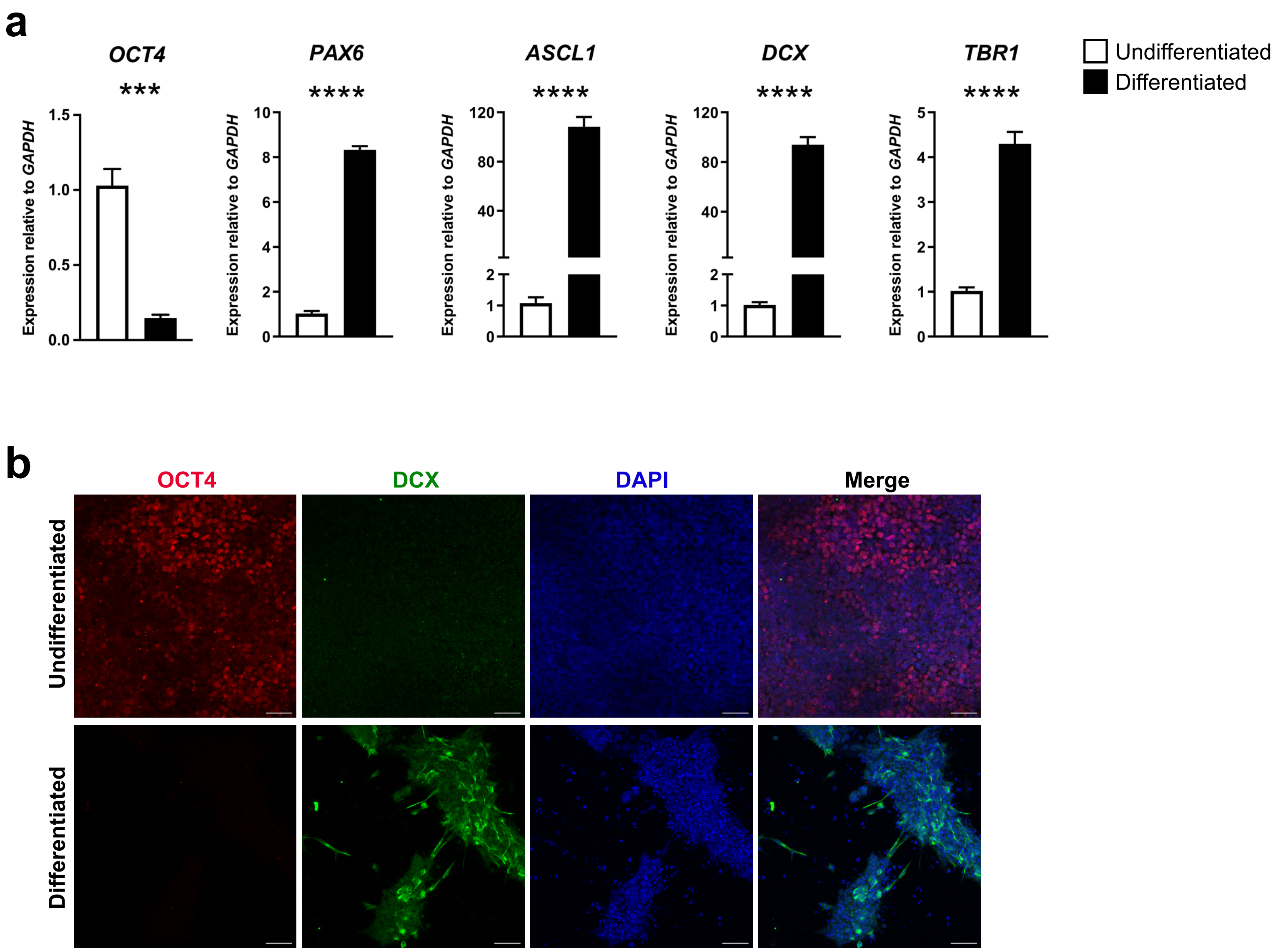

### Figure S2

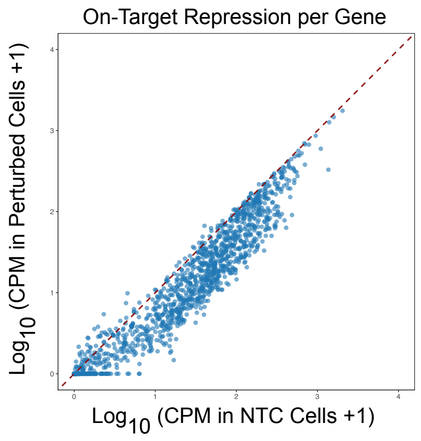

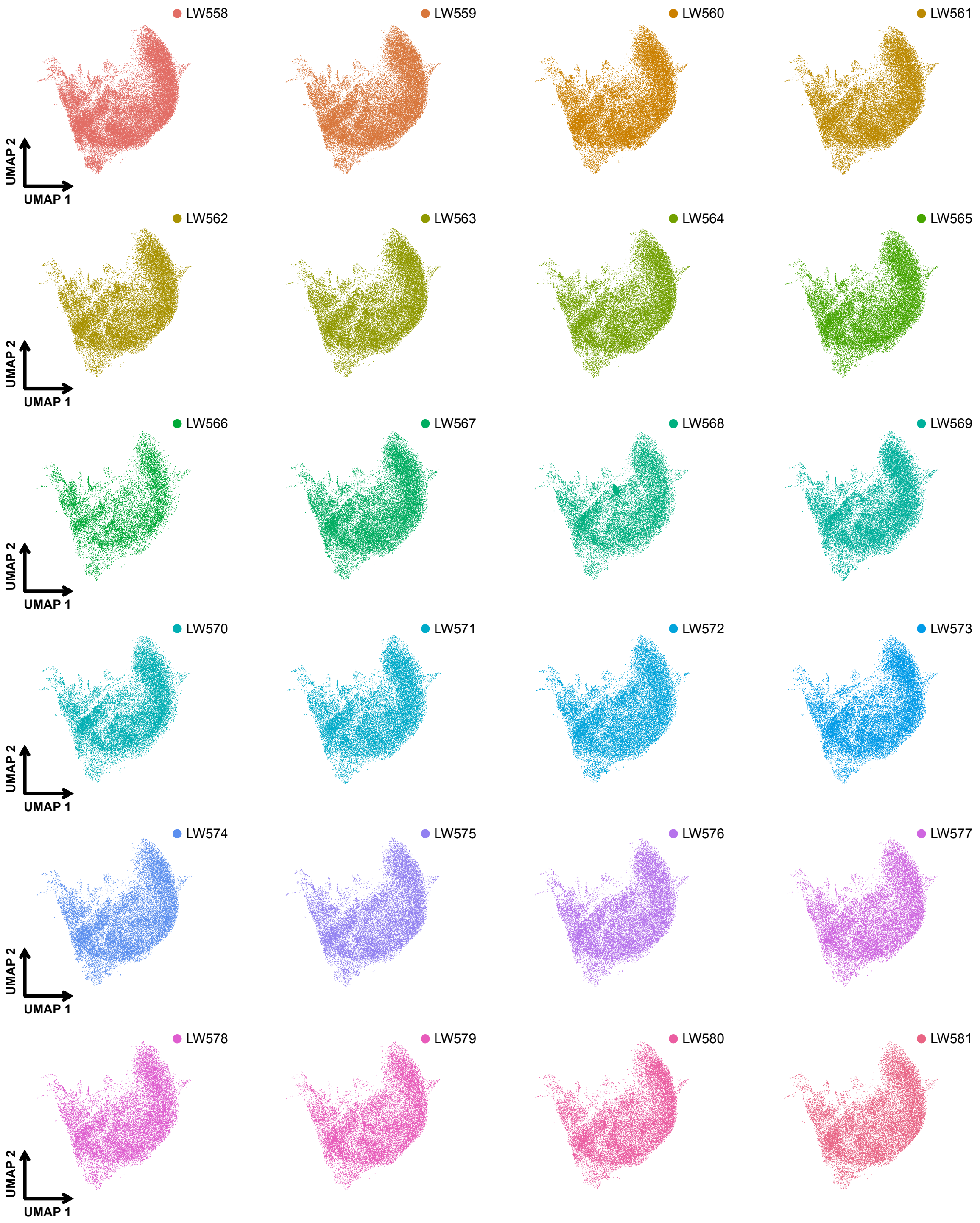

### Figure S3

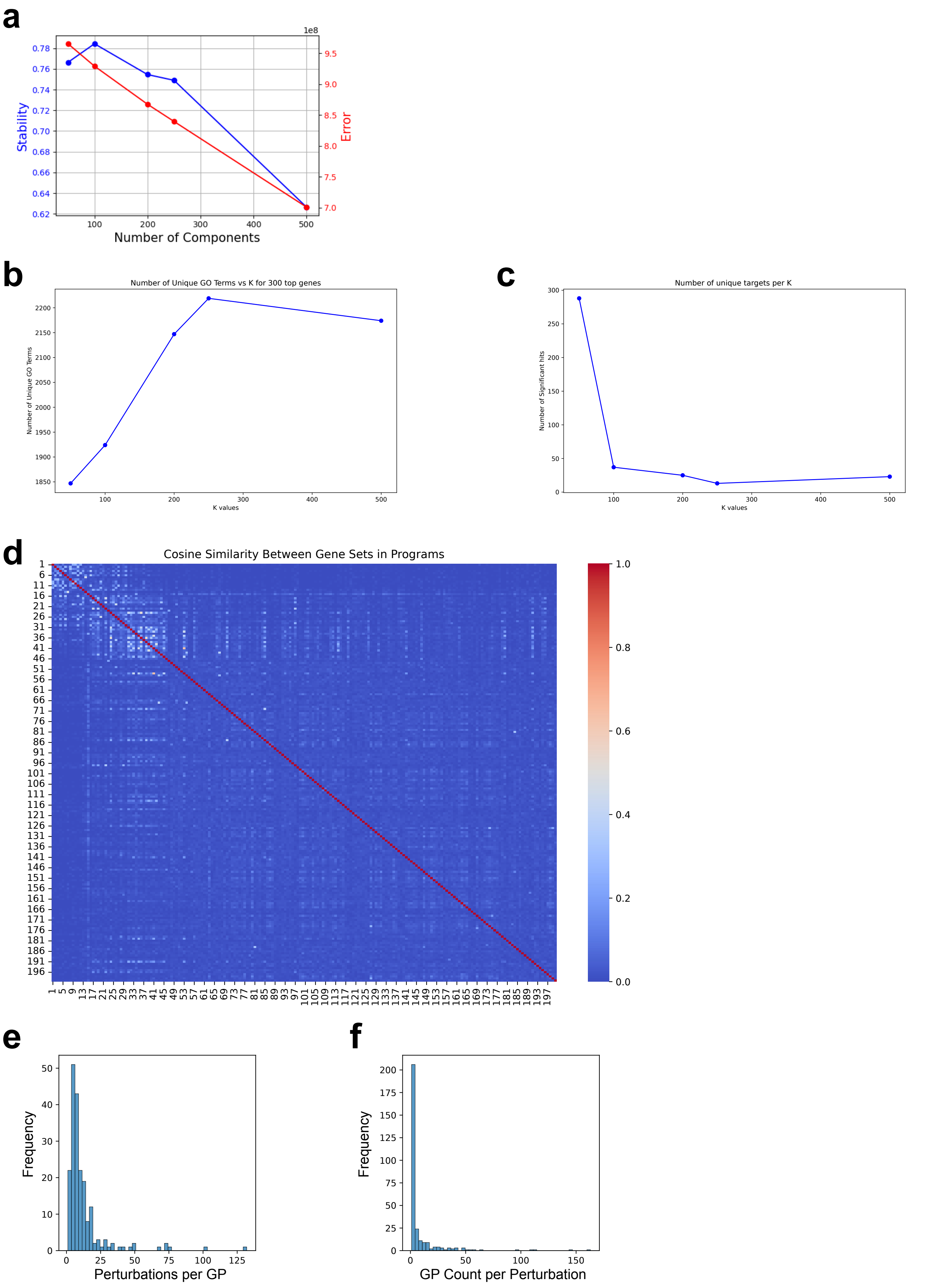

### Figure S4

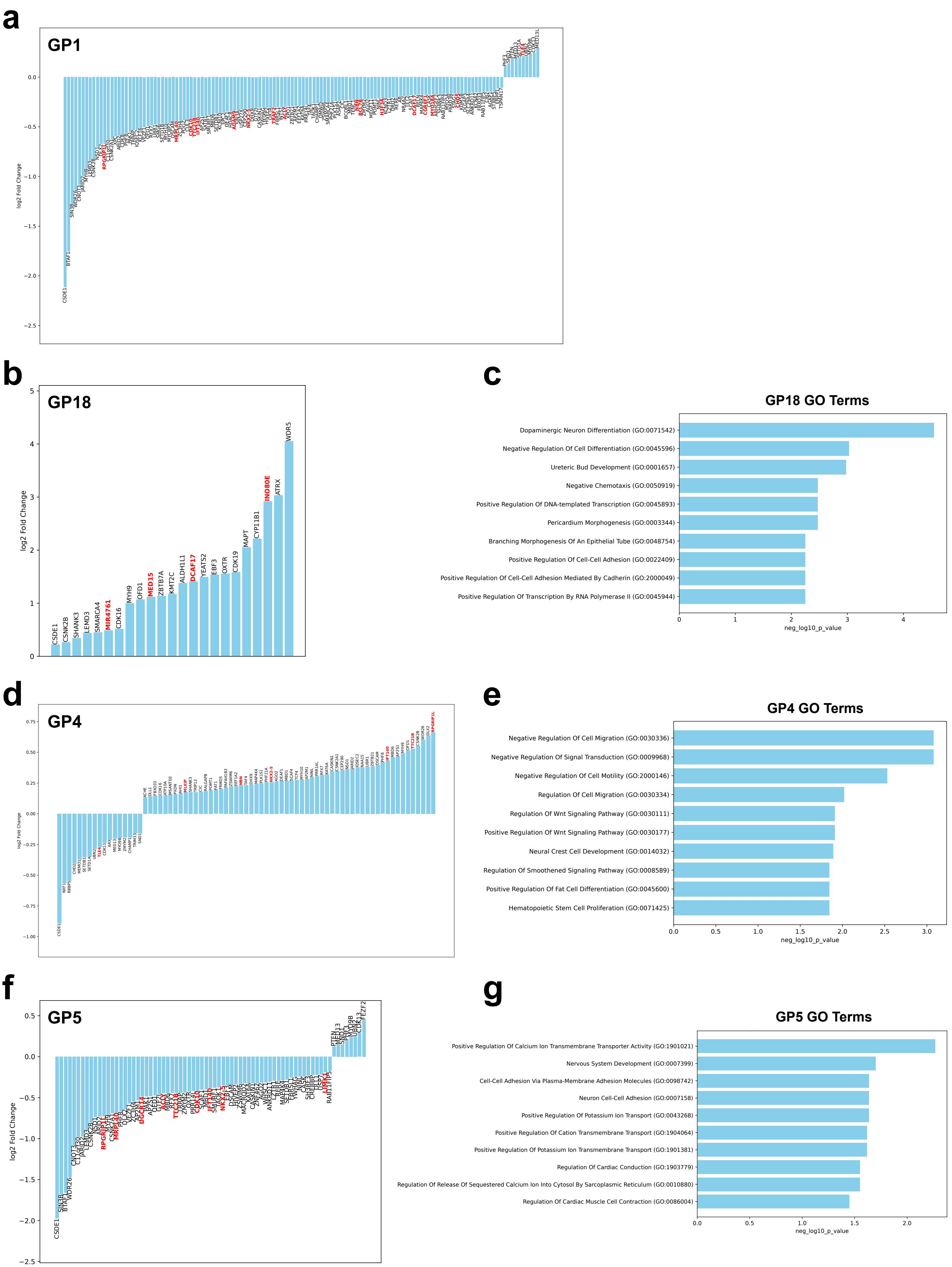

### Figure S5

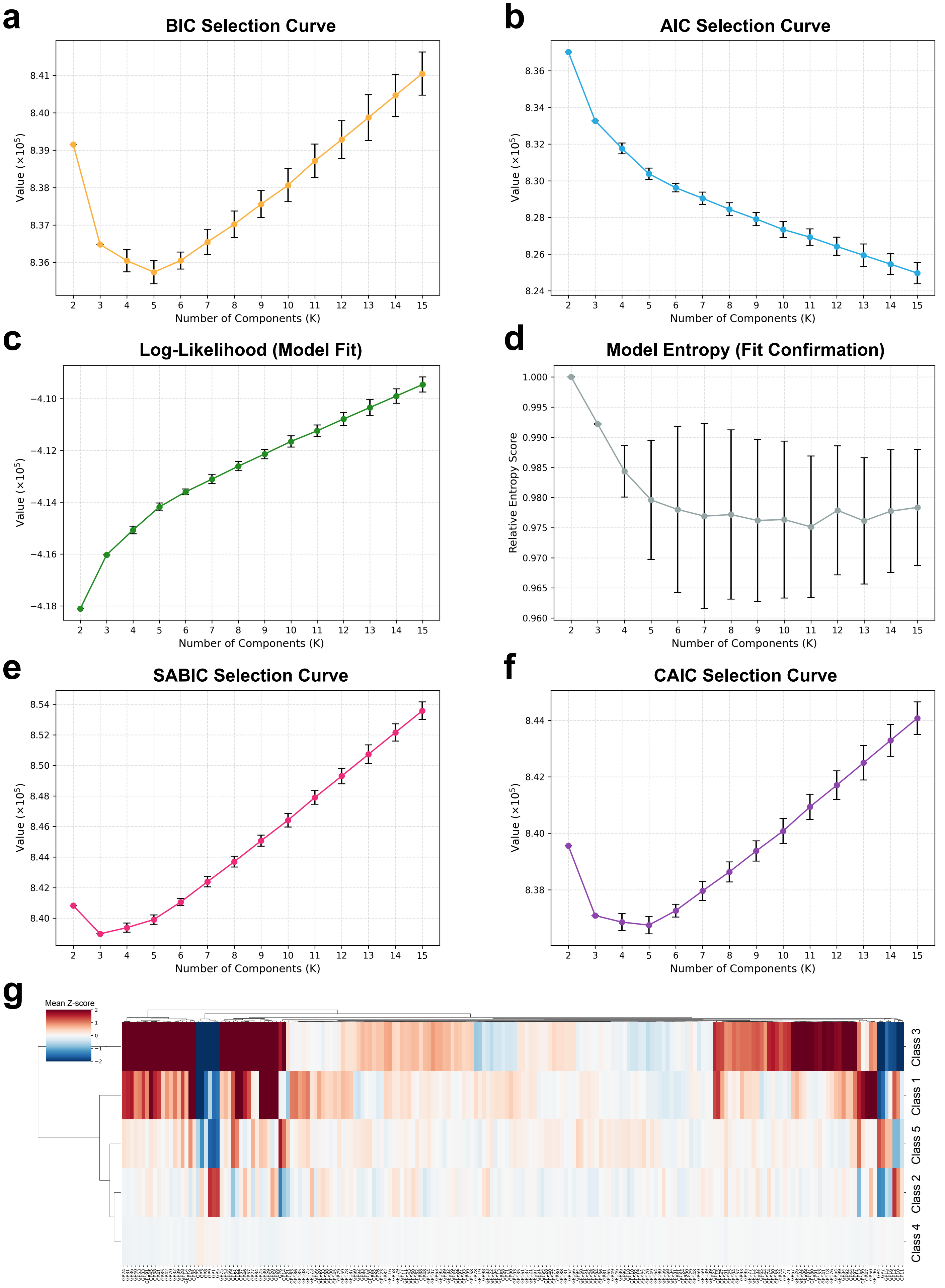

### Figure S6

#### Figure S6

**a**

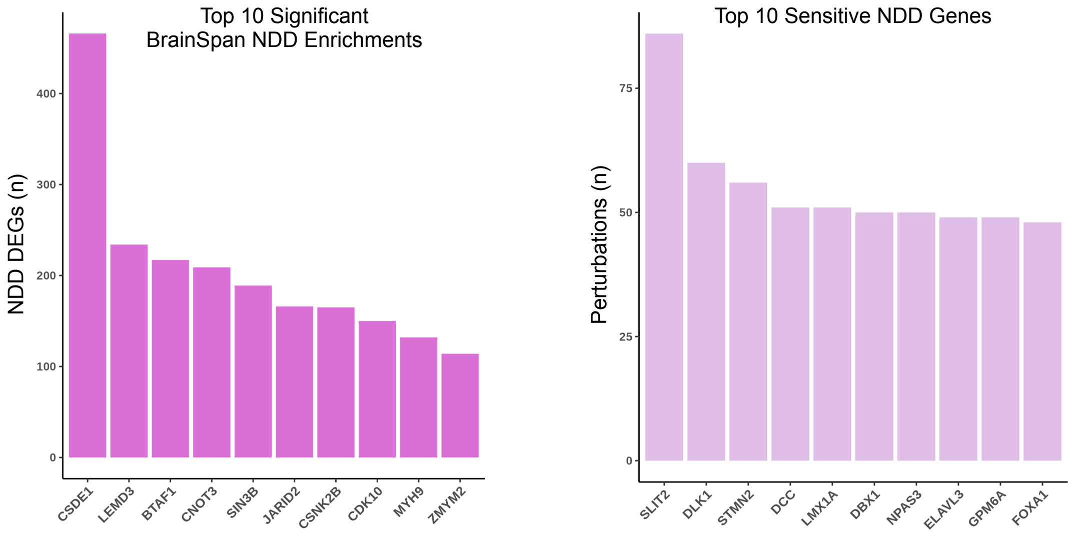


**b**

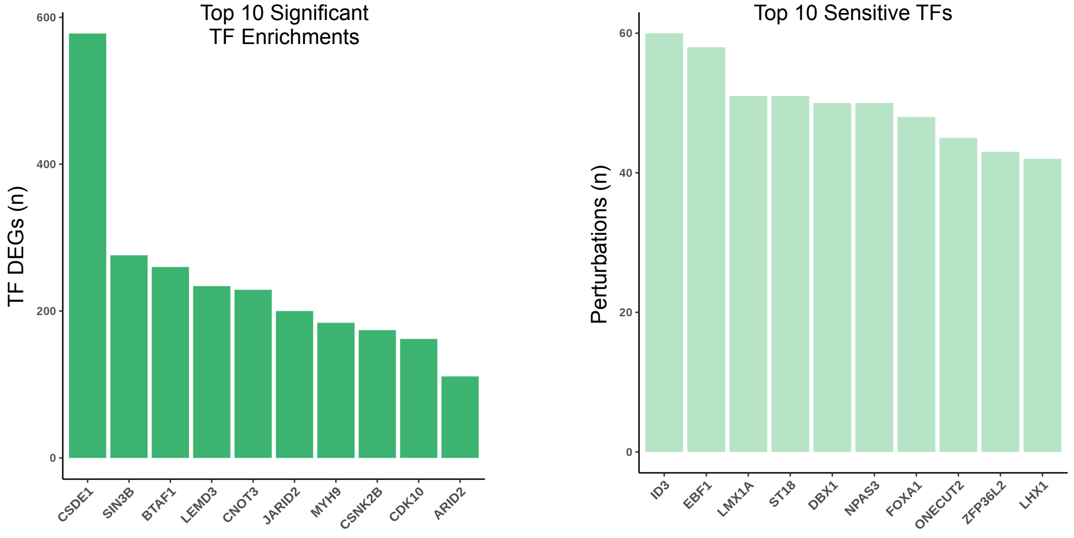


**c**

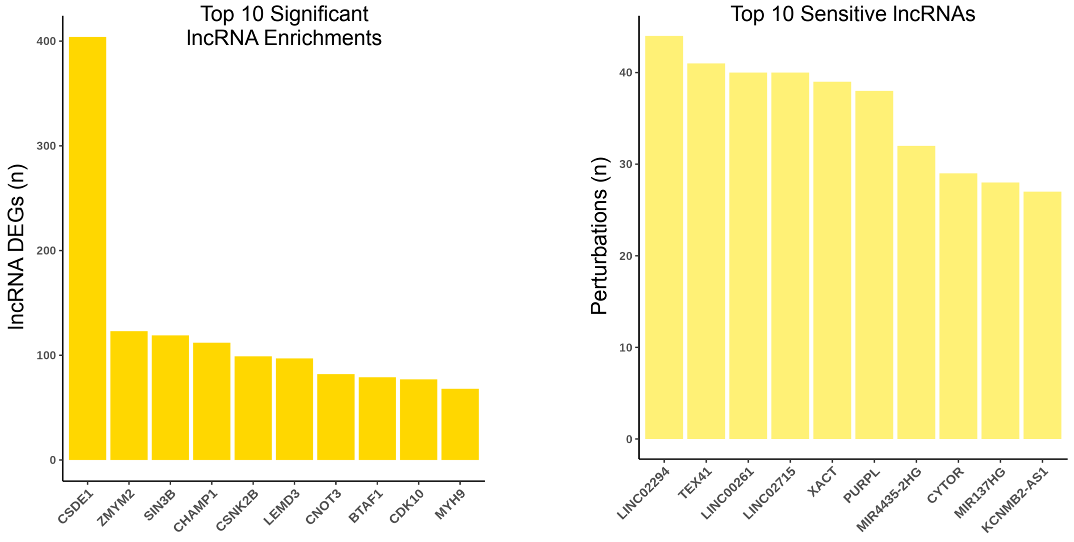
